# A nucleolar assembly module integrates an ancestral isoaspartylase to safeguard ribosome biogenesis

**DOI:** 10.64898/2026.09.04.749427

**Authors:** Alexander Gregor Geiger, Sébastien Favre, Oscar Vadas, Tobias von Arx, Fabian Ackle, Michael Alexander Ruoss, Liridona Zyberaj, Yash Verma, Ryan Separovich, Cedric A. J. Hutter, Michaela Oborská-Oplová, Daniela Portugal-Calisto, Markus A. Seeger, Pedro Beltrao, Nicolas Leulliot, Alexandre Smirnov, Dieter Kressler, Vikram Govind Panse

## Abstract

Eukaryotes inherited the core ribosome biogenesis apparatus from archaea. However, nucleocytoplasmic compartmentalisation and expansion to >200 assembly factors created the challenge of integrating this ancestral machinery into a complex maturation programme. One solution is the formation of transient modules in which newly acquired assembly factors support deeply conserved components. Here, we identify such a module, centred on the ancestral isoaspartylase Fap7, which couples the modification of the ribosomal protein uS11 to its incorporation into pre-ribosomes. Fap7 partners with Krr1 to capture uS11, forming an early checkpoint in which uS11 loading licenses Kri1 engagement and assembly progression. Loss of uS11 modification triggers a late checkpoint, preventing aberrant pre-ribosomes from acquiring translational competence. Integrative structure-function studies reveal how the ancestral isoaspartylase is embedded within a conserved eukaryotic assembly-factor network to safeguard the timing, order, and fidelity of ribosome production.

## Introduction

The emergence of the nuclear envelope in archaeal ancestors uncoupled transcription from translation, enabling additional layers of gene regulation, RNA processing, and surveillance. This necessitated the reorganisation of the pre-existing machinery that operated within a single cytoplasmic environment. How ancestral pathways were rewired to coordinate processes across nuclear and cytoplasmic compartments remains poorly unexplored^1^.

Ribosome assembly, a conserved pathway that couples pre-rRNA synthesis with ribosomal protein (r-protein) incorporation, was amongst the several processes impacted during eukaryogenesis. In anucleate bacteria and archaea, ribosome assembly is streamlined by co-transcriptional association of nascent r-proteins with emerging pre-rRNA^2,3^. The entire process is assisted by a handful of ribonucleases, RNA helicases, modification enzymes, and GTPases^2–4^. Although eukaryotic ribosomes retain architectural features inherited from archaea, including conserved rRNA folds and core r-proteins^3^, their assembly pathway has diverged markedly in complexity. Unlike archaeal ribosome biogenesis, which involves few dedicated assembly factors, eukaryotic 40S and 60S subunit biogenesis depends on >200 conserved factors acting sequentially across the nucleolus, the nucleoplasm, and the cytoplasm^5–7^. In the eukaryotic model budding yeast, the assembly cascade is initiated in the nucleolus during the production of the 35S pre-rRNA transcript by RNA Pol-I^8^. The emerging transcript engages with ∼100 factors and numerous small nucleolar ribonucleoprotein (snoRNP) complexes to form the 90S pre-ribosome, the earliest 40S subunit precursor^9^. Endonucleolytic cleavage of the nascent transcript separates the assembly pathway of 40S and 60S subunit precursors, which transiently associate with numerous RNA helicases, AAA^+^-ATPases, GTPases, nucleases, and chaperones that remodel pre-rRNA, and drive pre-ribosomes towards export competence^10^. Following nuclear export, pre-ribosomes undergo final maturation in the cytoplasm, providing a time window to proofread their functionality prior to entering translation^5,11^. These quality control steps are intricately linked to the sequential release of the remaining associated assembly factors, incorporation of remaining r-proteins, and pre-rRNA trimming and processing^12,13^. For example, cytoplasmic maturation of the 40S pre-ribosome involves the cleavage of 20S pre-rRNA by the endonuclease Nob1 to generate mature 18S rRNA, an essential step that licenses the 40S subunit for translation^14–16^. Such a spatial separation of assembly and quality control steps necessitates coordination with cellular trafficking pathways.

Another constraint introduced by compartmentalization is the correct targeting of nascent r-proteins from the cytoplasm to the nucleolus for incorporation into pre-ribosomes. Before every cell division, a yeast nucleus imports ∼14 million r-proteins from the cytoplasm to the nucleus for ribosome assembly. r-proteins are small, highly basic, and non-native in their unbound state, rendering them unstable and aggregation-prone. To deal with this logistical challenge, eukaryotes employ a network of importins, escortins, and dedicated chaperones that bind to their r-protein clients and mediate their transfer to pre-ribosomes^5,17–19^. How these delivery factors interface with the assembly machinery in space and time to coordinate r-protein folding and incorporation into developing pre-ribosomes remains unclarified.

The integration of the r-protein uS11 (yeast Rps14) into the nucleolar localised 90S pre-ribosome exemplifies this coordination challenge. The process relies on the nuclear localised isoaspartylase Fap7, inherited by eukaryotes from their archaeal ancestor^20–22^. Fap7 binds to uS11 and catalyses the ATP-dependent isomerisation of a specific aspartate residue within its disordered C-terminal tail into an isoaspartate (isoAsp)^22^. This enzymatic maturation of uS11 is critical to assemble the rRNA platform in the proximity of the mRNA exit site and organise the decoding centre on the 40S subunit. It is unclear how uS11 isoaspartylation is synchronised with its incorporation into the 40S pre-ribosome and alongside ongoing maturation steps. Here, we combined proximity-dependent biotinylation, hydrogen-deuterium exchange mass spectrometry, structural and genetic studies in yeast to unveil the functional environment of Fap7, which has remained refractory to cryo-EM studies. We show that the ancestral Fap7 fold has structurally evolved to embed its essential isoaspartylase activity into an intricate nucleolar assembly-factor network and guarantee spatiotemporal coordination of eukaryotic ribosome biogenesis across cellular compartments.

## Results

### Fap7 mediates conformational maturation of uS11 during 40S assembly

The r-protein uS11 occupies and organizes the platform of the 40S subunit, where it bridges conserved 18S rRNA helices essential for translation^23,24^. Although universally conserved, mechanisms that assemble uS11 into pre-ribosomes have diverged during evolution. In bacteria (and in mitochondria), uS11 integration into the pre-ribosome depends on YbeY (Fig. 1a)^25–29^. In archaea and eukaryotes, this process necessitates the ATPase Fap7 (Fig. 1a)^20,21,30,31^. Despite the lack of structural homology, both YbeY and Fap7 harbour isoaspartylation activity that modifies a specific asparagine and aspartate residue, respectively, within uS11 to an isoAsp^22^.

**Fig. 1.**
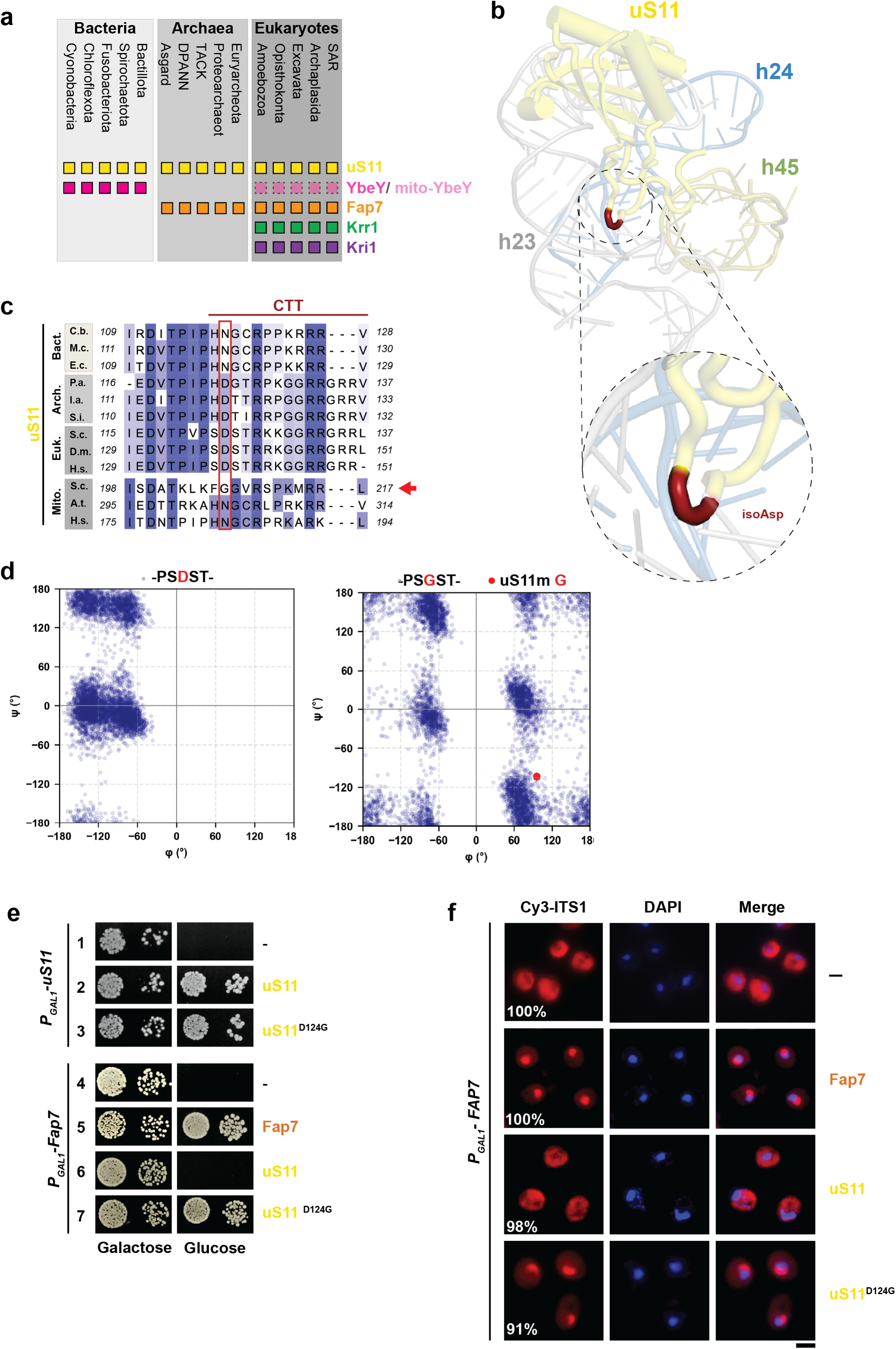
The essential role of Fap7 is conformational maturation of uS11^CTT^. **a**, Evolutionary distribution of r-protein uS11 and the assembly factors Fap7, Krr1, and Kri1 across phylogenetic groups. Squares indicate the presence of the corresponding protein in representative lineages from the three domains of life: Eukaryotes, Archaea, and Bacteria. **b**, Cryo-EM structure of a 40S pre-ribosome (PDB: 6RBD) showing uS11 (yellow) bound to 18S rRNA adjacent to helices h23 (grey), h24 (light blue), and h45 (orange). The isoAsp within uS11^CTT^ is highlighted in red. **c,** Multiple sequence alignment of the uS11^CTT^ from representative bacteria (Bact.), archaea (Arch.), eukaryotes (Euk.), and mitochondrial protein variants (Mito.). Amino acid conservation is shown in shades of blue, and mutated residues are highlighted by red boxes. Sequences are from *Cyanobacteria* (C.b.), *Microcystis* (M.c.), *Escherichia coli* (E.c.), *P. abyssi* (P.a.), *I. aggregans* (I.a.), *S. islandicus* (S.i.), *S. cerevisiae* (S.c.), *A. thaliana* (A.t.), and *H. sapiens* (H.s.). The alignment was generated with COBALT^93^ and visualized with Jalview^84^. **d**, Ramachandran maps showing the conformational sampling of the aspartate or glycine, highlighted in red, within the depicted peptides. Each blue dot represents the φ and ψ backbone dihedral angles of this residue from an individual frame of the molecular dynamic’s simulation. The red dot indicates the φ and ψ angles of the Gly residue in uS11m within the yeast mito-ribosome (PDB: 5MRC). **e**, Growth analysis of the *P*_GAL1_-*uS11* and *P*_GAL1_-*FAP7* strains containing empty vector (-) or plasmids expressing Fap7, uS11 or uS11^D124G^. Cells were spotted in serial tenfold dilutions on a control galactose-containing medium (left) and on the Fap7 and uS11-depleting glucose-containing medium (right) and incubated at 30°C for 4 days. **f,** Localization of 20S pre-rRNA by FISH using a Cy3-labelled oligonucleotide complementary to the 5′ region of ITS1 (red). Nuclear and mitochondrial DNA were stained with DAPI (blue). A *P_GAL1_*-*FAP7* strain containing an empty vector (-) or plasmids expressing Fap7, uS11 or uS11^D124G^ were grown to mid-log phase in Fap7-depleting glucose containing minimal media at 25°C prior to processing the cells for imaging. Images were processed using ImageJ version 1.54. Percentage indicates the penetrance of the depicted phenotype. Scale bar, 5 μm.

The isoAsp modification increases backbone flexibility, allowing access to otherwise inaccessible conformations^32,33^. We hypothesised that this modification enables a characteristic tight kink (Fig 1b, inset), permitting the productive engagement of uS11 C-terminal tail (uS11^CTT^) with rRNA helices h23, h24, and h45 to assemble the platform region (Fig. 1b). Budding yeast mitochondria are one of rare biological systems that do not isoaspartylate their r-protein, uS11m. Instead, the corresponding position in uS11m is occupied by glycine (Fig. 1c, red arrow). This raised the possibility that glycine may provide an alternative structural solution that is otherwise fulfilled by isoAsp. To test this in silico, we performed molecular dynamics simulations on two peptides derived from uS11 (Fig 1c, Euk.), wild-type PS<u>D</u>ST and mutant PS<u>G</u>ST. Backbone φ and ψ dihedral angles were extracted for the central aspartate or glycine residue and visualized on Ramachandran maps (Fig. 1d). The aspartate-containing peptide displayed restricted φ/ψ distribution, with most conformations confined to negative φ values, indicating limited local backbone flexibility (Fig. 1d, left panel). In contrast, the glycine substitution broadened the conformational space sampled by the peptide (Fig. 1d, right panel), including positive φ conformations that were poorly populated by the aspartate variant. Notably, the φ and ψ angles of the corresponding glycine in uS11m within the yeast mitoribosome structure (PDB: 5MRC) fall within the conformational space uniquely sampled by the glycine-containing peptide (Fig. 1d, right panel, red dot). Thus, replacing aspartate with glycine in the peptide allows its backbone to access conformations comparable with those observed for yeast mitochondrial uS11m.

We investigated whether the aspartate-to-glycine substitution in yeast uS11 (uS11^D124G^) retained uS11 function. uS11^D124G^ was functional, as it complemented the lethality associated with uS11 depletion (Fig. 1e, rows 1-3). We tested whether the D124G substitution bypassed the essential requirement of Fap7 in vivo. This was indeed the case: expression of uS11^D124G^, but not uS11, expressed from a centromeric plasmid, rescued the lethality associated with Fap7-depleted cells in a dominant manner (Fig. 1e, rows 4-7). A characteristic phenotype of Fap7 depletion in yeast is the cytoplasmic accumulation of 40S pre-ribosomes that contain an Internal <u>T</u>ranscribed <u>S</u>pacer (ITS1) RNA, consistent with defective Nob1-mediated cleavage of 20S pre-rRNA^20,21^. Expression of uS11^D124G^, but not uS11, restored 20S pre-rRNA processing, in a dominant manner, as assessed by fluorescence in situ hybridization (FISH) using a Cy3-labelled DNA probe that base pairs with ITS1-RNA (Fig. 1f). These functional studies establish Fap7 as a dedicated conformational maturation factor for uS11. We conclude that Fap7-catalysed uS11 modification is essential for exported 40S pre-ribosomes to achieve translation competence.

### TurboID reveals Fap7 proximity to nucleolar localised assembly factors

The spatiotemporal context of Fap7-mediated uS11 isoaspartylation in vivo as well as how this modification is coupled to ribosome maturation remained unclear. We applied proximity-dependent biotinylation to define the functional environment of Fap7 in yeast. This approach employs a promiscuous biotin ligase fused to a protein of interest to covalently label neighbouring proteins in its spatial proximity. Labelled proteins can be efficiently enriched by streptavidin affinity capture and identified by mass spectrometry. We used the enhanced ligase TurboID, which is well suited to detect transient interactions in yeast^34–36^. N- or C-terminal TurboID fusions of Fap7 were conditionally expressed from a copper-inducible promoter. Labelled proteins were enriched under stringent conditions and analysed by label-free quantitative mass spectrometry. As controls, we employed cytoplasmic and nuclear localised TurboID-tagged yEGFP fusion proteins to determine background labelling in the two cellular compartments^36^. As expected, both TurboID-fused Fap7 baits enriched uS11, consistent with its role as a Fap7 client (Fig. 2a, yellow). The other prominent protein enriched in both datasets was the 90S pre-ribosome associated assembly factor Krr1 (Fig. 2a, green), which stabilises the central domain of the 20S pre-rRNA^37^. Cryo-EM studies show how Krr1 binds to uS11 near rRNA helix h23 (Fig. 2b, inset), supporting observed genetic interactions between the two proteins^38–44^. In addition, Kri1, an interaction partner of Krr1^41^, also enriched in both datasets (Fig. 2a, purple). Further other enriched factors included Faf1 in the N-terminal TurboID dataset (Fig. 2a, left panel; Fig. 2b, inset) and Imp3 - an interactor of Faf1 located next to Krr1 on the 90S pre-ribosome (Fig. 2b, inset) - in the C-terminal TurboID dataset (Fig. 2a, right panel). Enrichment of these conserved nucleolar factors was selective, since we did not find snR30 snoRNP-associated proteins that are present in the vicinity of Krr1 on another early state of the 90S pre-ribosome (Fig. 2c)^41^. Likewise, no enrichment of other assembly factors present on 90S / 40S pre-ribosomes, such as Pno1, Kre33, Enp2 and Rrp9, or the late-associating Rio2 could be observed (Fig. 2a).

**Fig. 2.**
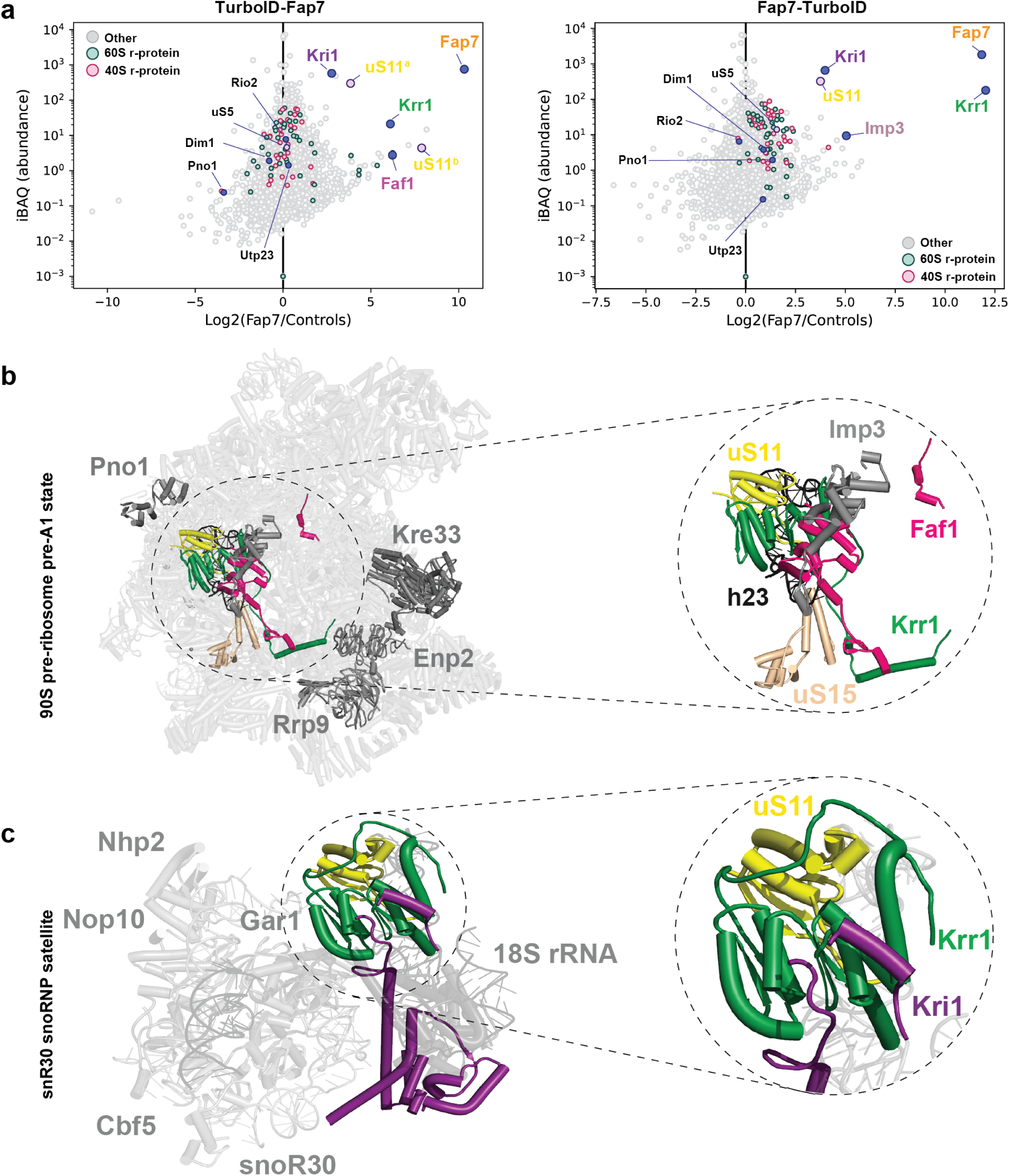
TurboID reveals the functional environment of yeast Fap7. **a**, TurboID-based proximity-labelling analysis of Fap7. TurboID was fused to either the N- or C-terminus of Fap7 (TurboID-Fap7 and Fap7-TurboID, respectively), and their local protein environment was analysed by mass spectrometry. Enriched proteins are highlighted in colour. Unique peptides corresponding to the two isoforms of uS11 (in yellow), uS11a and uS11b were detected in the TurboID-Fap7 assay. For each detected protein, normalized abundance, measured by intensity-based absolute quantification (iBAQ), is plotted against relative abundance, shown as log₂-transformed enrichment. Relative abundance was calculated by comparison with the mean protein abundance in two control purifications from cells expressing TurboID-GFP and NLS-TurboID-GFP (for TurboID-Fap7) or GFP-TurboID and NLS-GFP-TurboID (for Fap7-TurboID). TurboID assays were performed in single experimental replicates. **b,** Cryo-EM structure of the 90S pre-ribosome in transition state A1 (PDB: 6LQP). **c**, Cryo-EM structure of the snR30 snoRNP satellite associated with the 90S pre-ribosome (PDB: 9G25).

We conclude that the molecular environment of Fap7 in vivo is characterised by its proximity to uS11 and nucleolar assembly factors centred around Krr1, reflecting its transient interactions with the 90S pre-ribosome.

### Adaptation of the ancestral Fap7 fold enables Krr1 engagement

Proximity-dependent biotinylation studies in vivo identified Krr1 as a prominent candidate in the vicinity of Fap7 (Fig. 2a). An interaction between the *Chaetomium thermophilum* (Ct) CtFap7 and CtKrr1 was previously detected in yeast two-hybrid (Y2H) assays^37,45^, and the co-expression and purification of CtFap7 and CtKrr1 in yeast resulted in co-enrichment of yeast uS11^37^. Whether Fap7 directly binds to Krr1, or whether these components only assemble into a ternary complex with uS11 was not addressed.

Krr1 harbours two KH domains followed by a long, disordered C-terminal tail (CTT) (residues 193-316; Fig. 3a, upper panel). Truncation studies showed that the Krr1^CTT^ alone binds to Fap7 in Y2H assays (Fig. 3a, lower panel, row 2). In vitro pull-down assays corroborated a direct physical interaction between Krr1^CTT^ and Fap7. GST-Fap7 bound to Krr1^CTT^ (Fig. 3b, lane 4), whereas GST-Tsr4, the dedicated chaperone for uS5 (yeast Rps2)^46–48^, did not (Fig. 3b, lane 2), confirming binding specificity. Progressive truncations indicated that residues 246-256, which encompass a conserved proline-rich patch (PRP) (Fig. 3c), contribute to the interaction with Fap7 (Fig. 3a, rows 3-5). To define the structural basis of this interaction, we determined a crystal structure of CtFap7 bound to Krr1^PRP^. For this, CtFap7 was fused to a PRP consensus (KEYTPFPPPQ) (Fig. 3c) via a three residue GSG linker to the C-terminus. Crystals formed readily within a few days and diffracted to 1.8 Å resolution (Supplementary Table 1). The asymmetric unit revealed two CtFap7-Krr1^PRP^ fusion molecules arranged in a bead on a chain configuration, in which the Krr1^PRP^ of one protomer engages the neighbouring Fap7 molecule within the crystal lattice (Fig. 3d, left panel). The structure shows how Krr1^PRP^ is wedged in a cavity formed between a β-strand and the C-terminal α-helix of CtFap7 (Fig. 3d, middle panel). The interaction is mediated by two contacts: F250 within Krr1^PRP^ inserts into a hydrophobic pocket of CtFap7 lined by V146, L148, I163, W166, and W170, constituting the principal anchoring interaction, while T248 is clamped between W166 and W170, also contributing to the positioning of Krr1^PRP^ within the groove (Fig. 3d, right panel). W166 and W170 define the binding cavity, stabilising the groove between the β-strand and the C-terminal α-helix to accommodate the Krr1^PRP^ sequence.

**Fig. 3.**
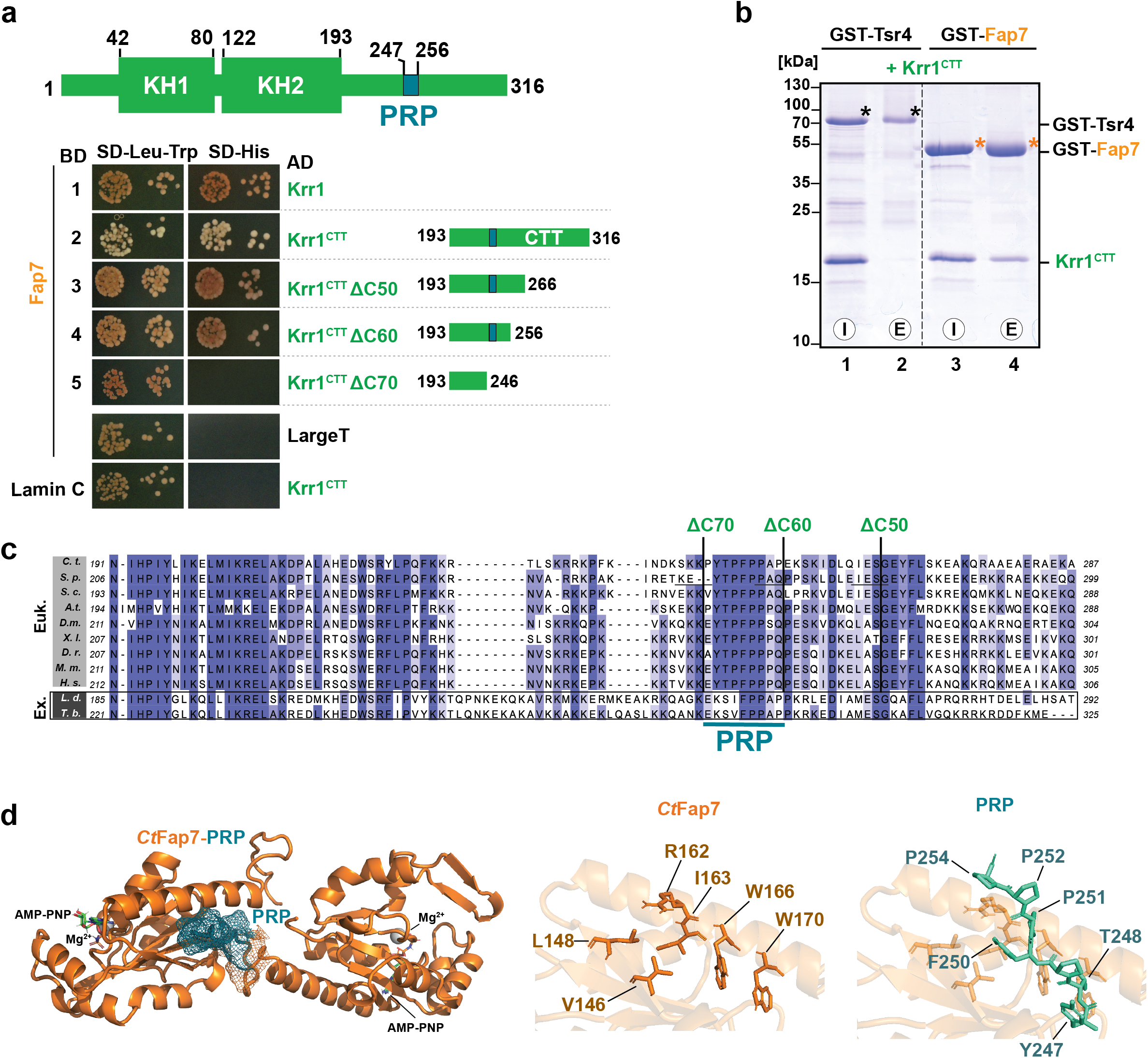
Krr1 interacts with Fap7 via a proline-rich patch (PRP). **a**, Top, domain organization of Krr1; the proline-rich patch (PRP) in the Krr1^CTT^ is shown in blue. Bottom, Y2H analysis of the Fap7-Krr1 interaction in NMY32 cells. The LexA DNA-binding domain (BD)-fused constructs are shown on the left and Gal4 activation domain (AD) fusions on the right. Transformants expressing the depicted fusions were spotted in serial tenfold dilutions on the indicated selective media and incubated at 30 °C for 4 days. Human lamin C (Lamin C) and SV40 large T antigen (Large T) served as negative controls. **b,** GST pull-down analysis of the interaction between Fap7 and the Krr1^CTT^. Recombinant GST-tagged Fap7 was immobilized on GSH-Sepharose beads and incubated with equimolar amounts of purified recombinant Krr1^CTT^. Beads were washed with PBS-Tween, and bound proteins were eluted with LDS sample buffer at 95°C, separated on a 4-20% Bis-Tris gradient gel, and visualized by Coomassie Blue staining. GST-tagged Tsr4 was used as a negative control. Asterisks indicate bait proteins. I, input; E, eluate. **c,** Multiple sequence alignment of the Krr1^CTT^ from representative eukaryotes (Euk.) and Euglenozoa (Ex.). Eukaryotic sequences are from *Chaetomium thermophilum* (C.t.), *Schizosaccharomyces pombe* (S.p.), *Saccharomyces cerevisiae* (S.c.), *Arabidopsis thaliana* (A.t.), *Drosophila melanogaster* (D.m.), *Xenopus laevis* (X.l.), *Danio rerio* (D.r.), *Mus musculus* (M.m.), and *Homo sapiens* (H.s.); Euglenozoa sequences are from *Leishmania donovani* (L.d.) and *Trypanosoma brucei* (T.b.). The alignment was generated with COBALT^93^ and visualized with Jalview^84^. **d,** Left, crystal structure of *C. thermophilum* Fap7 (CtFap7; orange) in complex with the CtKrr1^PRP^ (turquoise). AMP-PNP and Mg²⁺ in the active centre are indicated. The meshwork marks the peptide-binding site. Middle and Right panels, close-up views of CtFap7 and CtKrr1^PRP^ residues involved in Krr1:PRP complex formation, respectively.

To assess the functional relevance of the Fap7:Krr1^PRP^ interaction in vivo, we sought to identify Fap7 variants that preserve the key interaction with uS11 but were perturbed in engaging Krr1. Sequence alignments show that archaeal Fap7 (aFap7) homologs lack the conserved tryptophan residues that line the Krr1^PRP^-binding site in eukaryotes (Fig. 4a). Structural comparisons between *Ct*Fap7 and aFap7 showed a high overall similarity but a marked divergence of the C-terminal region (Fig. 4a, 4b) ^30^. In the case of aFap7, an archaeal-specific loop (red), followed by an extra α-helix, sterically occludes the interaction interface with Krr1^PRP^. Moreover, aFap7 lacks the β-strand/α-helix cavity required to accommodate Krr1^PRP^ (Fig. 4b). This non-productive C-terminal configuration is conserved amongst archaeal Fap7 homologs and differs from eukaryotes (Fig. 4a, red box), suggesting that the C-terminal portion of the ancestral Fap7 fold has been remodelled during evolution to enable Krr1^PRP^ recruitment. Consistent with these observations, Y2H (Fig. 4c, compare rows 1 & 3) and in vitro pull-down assays (Fig. 4d, compare lanes 4 & 5) show that aFap7 does not bind to Krr1^CTT^, while retaining interactions with yeast uS11 (Fig. 4c, compare rows 2 & 4). The consequences of this uncoupling were evident in vivo. Although aFap7, like yeast Fap7, enriched in the nuclear compartment (Fig. 4e), it failed to rescue the lethality of Fap7 depletion (Fig. 4f) and did not restore associated 20S pre-rRNA processing defects as assessed by FISH experiments (Fig. 4g). These data indicate that the uS11-binding activity of aFap7 alone is insufficient to support its role during 40S subunit assembly. We suggest that remodelling of the ancestral Fap7 fold generated a Krr1-binding pocket to enable its integration into a conserved eukaryotic assembly-factor network.

**Fig. 4.**
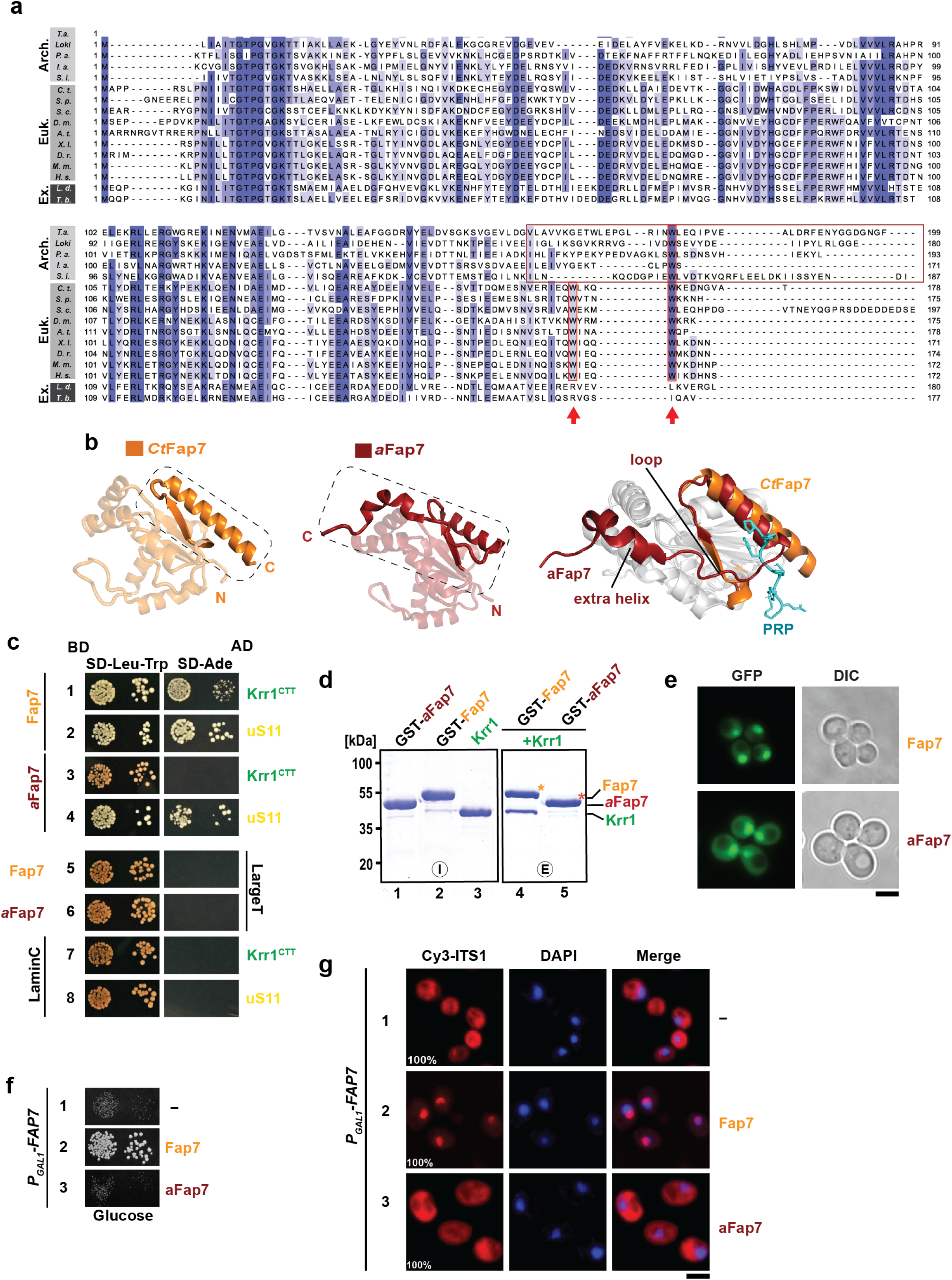
Eukaryotic features permit Fap7:Krr1 interactions. **a**, Multiple sequence alignment of Fap7 from representative archaea (Arch.), eukaryotes (Euk.), and Euglenozoa (Ex.). Archaeal sequences are from *Pyrococcus abyssi* (P.a.), *Lokiarchaeum ossiferum* (Loki.), *Ignisphaera aggregans* (I.a.), *Saccharolobus islandicus* (S.i.), *Thermosphaera aggregans* (T.a.); eukaryotic sequences are from *C. thermophilum* (C.t.), *S. pombe* (S.p.), *S. cerevisiae* (S.c.), *A. thaliana* (A.t.), *D. melanogaster* (D.m.), *X. laevis* (X.l.), *D. rerio* (D.r.), *M. musculus* (M.m.), and *H. sapiens* (H.s.); Euglenozoa sequences are from *L. donovani* (L.d.) and *T. brucei* (T.b.). The alignment was generated with COBALT^93^ and visualized with Jalview^84^. **b,** Left, crystal structure of CtFap7, with the C-terminal α-helix highlighted in a dashed box. Middle, crystal structure of archaeal *P. abyssi* Fap7 (aFap7; PDB: 4CVN^31^, with the C-terminal α-helices highlighted in a dashed box. Right, structural superposition of CtFap7 (orange) and *P. abyssi* aFap7 (red). The archaeal-specific α-helix and loop are indicated. **c,** Y2H analysis of yeast or archaeal Fap7 and yeast Krr1^CTT^ and uS11. Lamin C and Large T served as negative controls. **d,** GST pull-down analysis comparing the interaction of archaeal and yeast Fap7 with Krr1. Purified GST-tagged archaeal Fap7 (aFap7) or yeast Fap7 immobilized on GSH-Sepharose beads were incubated with equimolar amounts of recombinant Krr1. Beads were washed with a buffer containing 300 mM NaCl, and the bound proteins were eluted with LDS sample buffer, separated by SDS-PAGE, and visualized by Coomassie Blue staining. Asterisks indicate the bait proteins. I, input; E, eluate. **e,** Localization of GFP fusions of yeast Fap7 and aFap7 in logarithmically growing yeast cells cultured in medium containing 2% glucose, as monitored by fluorescence microscopy. Images were processed using ImageJ version 1.54. A representative image from *n*=3 biological replicates is shown. **f,** Growth analysis of a *P_GAL1_*-*FAP7* strain containing an empty vector (-) or plasmids expressing yeast Fap7 or aFap7 under control of the yeast *FAP7* promoter. Cells were spotted in serial tenfold dilutions on a control galactose-containing medium (left) and on the Fap7-depleting glucose-containing medium (right) and incubated at 30°C for 4 days. **g,** Localization of 20S pre-rRNA by fluorescence in situ hybridization (FISH) using a Cy3-labelled oligonucleotide complementary to the 5′-region of ITS1 (red). Nuclear and mitochondrial DNA were stained with DAPI (blue). The *P_GAL_*_1_-*FAP7* cells transformed with an empty vector (-) or plasmids expressing yeast Fap7 or aFap7 were grown in Fap7-depleting glucose containing medium to mid-log phase at 25°C. Percentage indicates the penetrance of the depicted phenotype. Scale bar, 5 μm.

The sequence divergence of the Fap7 C-terminal region within the early branching Euglenozoa relative to other eukaryotes drew our attention (Fig. 4a, see Ex.). While Fap7 homologues from *Leishmania donovani* and *Trypanosoma brucei* (Tb) retain the eukaryotic Fap7 fold (Fig. 5a, left panels) including the C-terminal region, they lack the tryptophan residues that are present in the Krr1^PRP^-binding pocket of opisthokonts (Fig 5a, right panels). Conversely, Krr1 sequences from these excavates harbour a slightly different PRP variant, KS[I/V]FPP as compared to the opisthokont YTPFPP consensus (Fig. 3c, see Ex.). Based on these analyses, we suspected that TbFap7 would exhibit an altered interaction with yeast Krr1. Indeed, we found that TbFap7 interacts with yeast Krr1, albeit weaker than yeast Fap7, as judged by the smaller colony size in Y2H assays (Fig. 5b, compare rows 1 & 2). Notably, TbFap7 did not interact with yeast Krr1^CTT^ alone in these assays (Fig. 5b, compare rows 3 & 4), but still showed robust binding to yeast uS11, consistent with its ancestral chaperoning activity (Fig. 5b, compare rows 5 & 6). Thus, TbFap7 represents a Fap7 variant that preserves uS11 binding while exhibiting a perturbed interaction with yeast Krr1. To assess the consequences of this attenuated interaction in vivo, we expressed TbFap7 from a centromeric plasmid in Fap7-depleted cells. While TbFap7 rescued the lethality of the Fap7 depletion, the resultant cells were growth-impaired (Fig. 5c, compare row 2 & 4) and accumulated 20S pre-rRNA in the cytoplasm as judged by FISH experiments (Fig. 5d). Over-expression of TbFap7 from a high-copy 2µ plasmid restored growth of the Fap7 depletion mutant (Fig. 5c, compare rows 2 & 5) and 20S pre-rRNA processing (Fig. 5d) close to wild-type levels. We infer that increased TbFap7 levels functionally compensate for the attenuated interaction with yeast Krr1 in vivo.

**Fig. 5.**
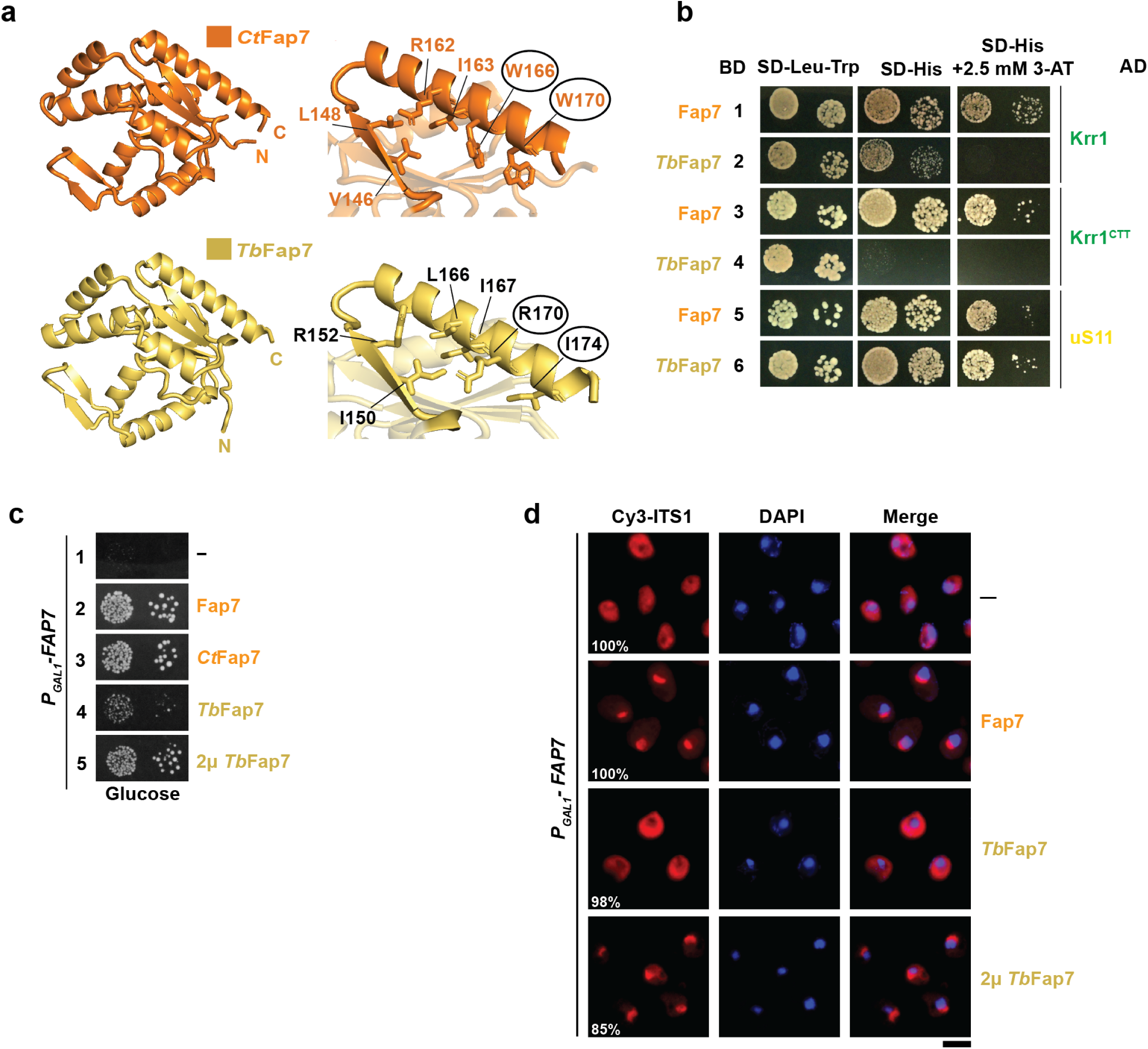
Functional complementation of *T. brucei* Fap7 in yeast. **a**, Top left, crystal structure of CtFap7. Bottom left, AlphaFold model of TbFap7. Top right, CtFap7 residues involved in Krr1^PRP^ binding are indicated, and two conserved tryptophan residues are highlighted with circles. Bottom right, the corresponding residues in TbFap7 predicted to contact Krr1^PRP^. **b,** Y2H analysis of the interactions between yeast Fap7 or *T. brucei* (TbFap7) with yeast Krr1, Krr1^CTT^ and uS11. The LexA DNA-binding domain (BD)-fused constructs are shown on the left and Gal4 activation domain (AD) fusions on the right. Transformants expressing the depicted fusions were spotted in serial tenfold dilutions on the indicated selective media and incubated at 30 °C for 4 days. **c,** Growth analysis of the *P*_GAL1_-*FAP7* strain containing empty vector (-), monocopy centromeric plasmids expressing yeast Fap7, CtFap7, or TbFap7 or a multicopy plasmid (2μ) expressing TbFap7. Cells were spotted in serial tenfold dilutions on a control galactose-containing medium (left) and on the Fap7-depleting glucose-containing medium (right) and incubated at 30°C for 4 days. **d,** Localization of 20S pre-rRNA by fluorescence in situ hybridization (FISH) using a Cy3-labelled oligonucleotide complementary to the 5′ region of ITS1 (red). Nuclear and mitochondrial DNA were stained with DAPI (blue). The *P_GAL1_*-*FAP7* strain was transformed with empty vector (-) or plasmids expressing yeast Fap7, centromeric TbFap7 or a high plasmid (2μ) expressing TbFap7. The transformants expressing the depicted plasmids were grown in Fap7-depleting minimal glucose containing medium to mid-log phase at 25°C. Micrographs were processed using ImageJ. Percentage indicates the penetrance of the depicted phenotype. Scale bar, 5 μm.

### uS11 allosterically modulates Krr1:Fap7 interactions

TbFap7 showed a weak interaction with yeast Krr1 and, accordingly, rescued the lethality of Fap7 depletion when overexpressed. We suspected that Krr1 contains additional interaction site(s) that reinforced TbFap7 binding. To delineate the interfaces, we reconstituted Krr1:Fap7 and Krr1:Fap7:uS11 complexes (Fig. 6a; 6b, left panels). Crystallization attempts were unsuccessful; therefore, we employed hydrogen-deuterium exchange mass spectrometry (HDX-MS) and AlphaFold modelling^49,50^ (Supplementary Figure 1a, 1b) to probe the architecture and dynamics of these complexes. Comparison of HDX rates between the target protein alone and when in a complex enables mapping of binding interfaces as well as allosteric changes^51–53^.

**Fig. 6.**
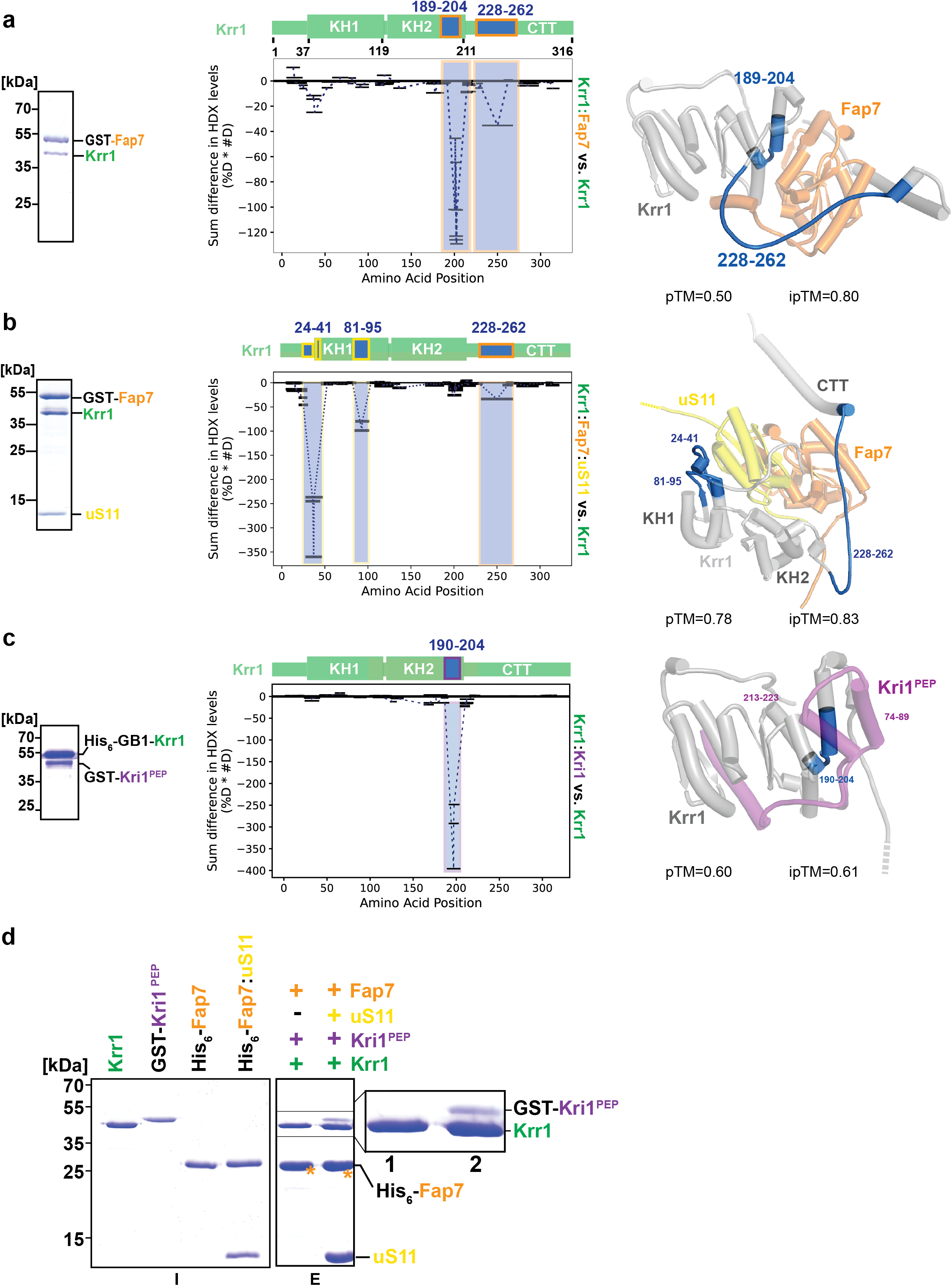
Organisation of Fap7:Krr1-containing complexes. **a-c**, Left panels, SDS-PAGE analyses of protein complexes purified from *E. coli* co-expressing yeast (a) GST-Fap7 and Krr1, (b) GST-Fap7, Krr1, and uS11 (b), or (c) GST-Kri1^PEP^ and His_6_-GB1-Krr1. Middle panels, HDX-MS analyses: the observed peptides are plotted according to their position in Krr1 (X-axis) as black lines and normalized deuterium uptake (Y-axis). Regions showing significant changes in HDX are highlighted within coloured boxes. Three technical replicates were performed. Source data are provided as Supplementary Data 2. Right panels, AlphaFold models showing protected regions of Krr1 (blue) in complex with Fap7, Fap7:uS11 or Kri1, as indicated^50^. **d,** His_6_-Ni^2+^-NTA pull-down studies of the Fap7:uS11:Krr1:Kri1^PEP^ complex. Recombinant His_6_-tagged Fap7 or the His_6_-Fap7:uS11 complex were incubated with equimolar amounts of untagged Krr1 and GST-Kri1^PEP^ with Ni^2+^-NTA beads. The immobilized complexes were washed prior to elution and analysed by SDS-PAGE. I, input; E, eluate. Asterisks indicate the His_6_-Fap7 bait.

To define how Krr1 interfaces with its interaction partners, we compared HDX rates of Krr1 alone and within reconstituted Krr1:GST-Fap7 and Krr1:GST-Fap7:uS11 complexes (Fig. 6a, 6b left panels). To this end, Krr1, Krr1:GST-Fap7, and Krr1:GST-Fap7:uS11 were exposed to D_2_O-containing buffer for times ranging from 3 s to 1 h, prior to quenching, proteolytic digestion and mass analysis of the peptides (Fig. 6; Supplementary Data 1). In the Krr1:GST-Fap7 complex, an extended region within the Krr1^CTT^ (residues 228-262), encompassing the PRP (residues 246-256), exhibited protection from exchange compared to isolated Krr1 (Fig. 6a, middle panel), corroborating our crystal structure (Fig. 3d). In addition, a second protected region spanning residues 189-204 within the KH2 domain was identified, in agreement with the AlphaFold model (Fig. 6a, middle & right panels). Strikingly, this protection of the KH2 region (residues 189-204) was lost in the context of the Krr1:GST-Fap7:uS11 complex (Fig. 6b, middle panel). Instead, new protected regions emerged within the Krr1 N-terminus (residues 24-41) and the KH1 domain (residues 81-95) (Fig 6b, middle panel). These Krr1 regions within the Krr1:Fap7:uS11 complex point to contacts between Krr1’s KH1 domain and uS11 and are supported by the AlphaFold model (Fig. 6b, right panel). Notably, the entire protected region within Krr1^CTT^ (residues 228-262) remained shielded within the ternary complex, indicating persistent engagement of this PRP-containing interface (Fig. 6a; 6b, middle panels). These findings indicate that Fap7 contacts Krr1 via the CTT (residues 228-262) and indirectly via uS11 through the Krr1 KH1 domain (Fig. 6b, middle and right panels). Further, they show that uS11 incorporation into the Krr1:Fap7 complex frees the Krr1 KH2 binding site, enabling interactions with an additional assembly factor (see below).

### uS11 licenses the Krr1:Fap7 complex to engage with Kri1

Another prominent factor identified in the Fap7 TurboID dataset was Kri1 (Fig. 2a, purple), an assembly factor that directly interacts with Krr1^42^. Kri1 functions during 90S pre-ribosome assembly, where it cooperates with Krr1 to anchor the snR30 snoRNP satellite (Fig. 2b, lower panel)^41^. Semi-quantitative proteomic analyses of 90S pre-ribosomes isolated via Utp10-FTpA (FTpA tag:3xFLAG-TEV-ProteinA) showed that recruitment of Kri1 and uS11 is strongly impaired upon Krr1 depletion^37^ (Supplementary Figure 2, left panel). Likewise, we found that uS11 depletion compromised Kri1 recruitment to 90S pre-ribosomes isolated via Krr1-FTpA (Supplementary Figure 2, right panel). Together, these proteomic datasets reveal a co-dependency for incorporation of Krr1, uS11, and Kri1 into the 90S pre-ribosome. To gain insights into the Krr1:Kri1 interface, we modelled the complex using AlphaFold2. The model shows how the three α-helices within the N-terminal region of Kri1 (residues 1-300) engage with the two KH domains of Krr1, with prominent contacts to the KH2 domain (Supplementary Figure 1c). Full-length Kri1 proved refractory to expression in *E. coli*. Hence, we engineered a construct, comprising residues 51-96 fused to residues 209-267 within the N-terminal region, hereafter referred to as Kri1^PEP^. Krr1 and Kri1^PEP^ co-expression in *E. coli* followed by pull-down studies confirmed the formation of a stable Krr1:Kri1^PEP^ complex (Fig. 6c, left panel).

We applied HDX-MS to validate the Krr1:Kri^PEP^ interaction surface (Supplementary Data 2). For this, we compared the HDX rates of Krr1 alone and within the Krr1:Kri^PEP^ complex. Binding of Kri1^PEP^ induced strong protection to HDX within residues 190-204 of the Krr1 KH2 domain, immediately preceding the CTT (Fig. 6c, middle panel). No such protection was observed for the KH1 domain. Consistent with the AlphaFold2 model (Fig. 6c, right panel), this protected region overlaps with the KH2 surface engaged by Fap7 within the Krr1:Fap7 complex (Fig. 6a, middle panel), which becomes only accessible upon binding to uS11. In agreement with these HDX-MS data, pulldown assays show that Kri1^PEP^ associates only with the ternary Krr1:Fap7:uS11 complex, but not with the binary Krr1:Fap7 complex, demonstrating the dependency of this interaction on uS11 (Fig. 6d, compare lanes 1 & 2, inset). We propose that uS11 loading relieves the steric clash of the KH2 domain within the ternary complex permitting interactions with Kri1.

### Krr1^CTT^ provides a versatile interaction platform for 90S pre-ribosome assembly

Our structural data show how that the C-terminal region of Fap7 has evolved in eukaryotes to interact with Krr1^PRP^. To examine this interaction from the reciprocal perspective, we investigated the functional contribution of the Krr1^PRP^ in vivo. Consistent with a previous report^41^, a 70-residue truncation of Krr1 (Krr1ΔC70) encompassing PRP failed to rescue the lethality of the Krr1-depletion mutant (Fig. 7a, compare rows 2 & 3), indicating that the C-terminal region is essential for Krr1 function. But a Krr1 variant with a mutated PRP (YTPFPP to SGGSGG; termed Krr1*) complemented the lethality of the Krr1 depletion mutant (Fig. 7a, compare rows 1 & 4). Accordingly, Y2H assays showed that Fap7 interacted with Krr1* to a similar extent as wild-type Krr1 (Fig. 7b, compare row 1 & 2). These results agree with HDX-MS data and the AlphaFold model, which indicate a broader protection of Krr1^CTT^ and interactions beyond the PRP (residues 246-256) (Fig 6a). We therefore generated a series of C-terminal truncations of Krr1 and Krr1* and tested their interactions with Fap7 using Y2H assays. We found that a 50-residue C-terminal truncation of Krr1, Krr1ΔC50, retained Fap7 binding, but the mutant, Krr1*ΔC50, only weakly interacted with Fap7 (Fig. 7b, compare rows 3 & 4). The Krr1*ΔC50 double mutant protein is not simply unstable since it robustly interacted with Faf1 (Fig. 7b, row 8), another 90S pre-ribosome associated assembly factor^54,55^ that enriched in Fap7 TurboID assays (Fig. 2a, left panel), indicating a specific impairment in Krr1:Fap7 contacts. Complementation studies showed that yeast cells expressing individual Krr1ΔC50 and Krr1* mutants were viable (Fig. 7a, rows 4 & 5), but the combined Krr1*ΔC50 double mutant was lethal (Fig. 7a, row 6). These data argue that multiple sites within Krr1^CTT^ cooperate to support its essential function(s) in vivo.

**Fig. 7.**
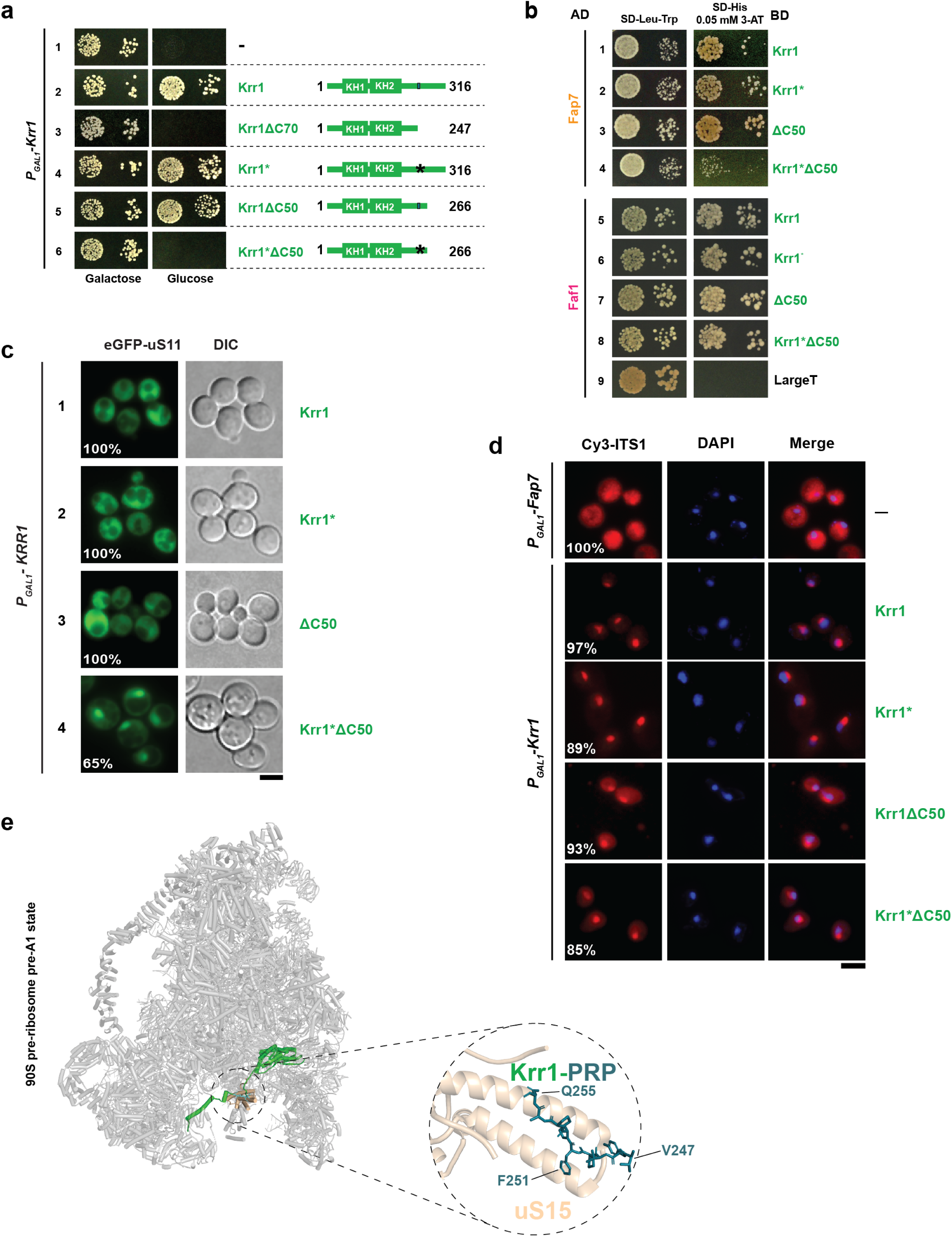
Multiple roles for Krr1^CTT^ during 40S pre-ribosome maturation. **a**, Growth analysis of the *P_GAL1_*-*KRR1* strain containing empty vector (-) or plasmids expressing Krr1, truncated Krr1ΔC70 and Krr1ΔC50, or their PRP-mutated counterparts Krr1*, truncated Krr1*ΔC50 variants. Cells were spotted in serial tenfold dilutions on a control galactose-containing medium (left) and on the Krr1-depleting glucose-containing medium (right) and incubated at 30°C for 4 days. **b,** Y2H analysis of the interactions between yeast Fap7 and Krr1 variants. The LexA DNA-binding domain (BD)-fused constructs are shown on the right and Gal4 activation domain (AD) fusions on the left. Transformants expressing the depicted fusions were spotted in serial tenfold dilutions on the indicated selective media and incubated at 30 °C for 4 days. Faf1 and Large T served as positive and negative controls, respectively for the experiment. **c,** Location of eGFP-uS11 in *P_GAL1_*-*KRR1* cells expressing Krr1, Krr1ΔC50, Krr1*, and Krr1*ΔC50. Cells expressing the indicated constructs and eGFP-uS11 were grown in Krr1-depleting minimal glucose containing medium to mid-log phase prior to imaging. Images were processed using ImageJ version 1.54. Percentage indicates the penetrance of the depicted phenotype. Scale bar, 5 μm. Representative images from *n*=3 biological replicates are shown. **d,** Localization of 20S pre-rRNA by fluorescence in situ hybridization (FISH) using a Cy3-labelled oligonucleotide complementary to the 5′ region of ITS1 (red). Nuclear and mitochondrial DNA were stained with DAPI (blue). The *P_GAL1_*-*KRR1* strain expressing Krr1, Krr1ΔC50, Krr1*, and Krr1*ΔC50 was grown in Krr1-depleting minimal glucose containing medium to mid-log phase at 25°C. The *P_GAL1_*-*FAP7* strain containing an empty vector (-) and grown on Fap7-depleting minimal miminal glucose containing media was used as a control for cytoplasmic 20S pre-rRNA accumulation. Images were processed using ImageJ version 1.54. Percentage indicates the penetrance of the depicted phenotype. Scale bar, 5 μm. **e,** Yeast Krr1^PRP^ bound to r-protein uS15 (yeast Rps13) within the 90S pre-ribosome pre-A1 state (PDB: 6LQP).

To investigate the functional significance of this synergistic effect, we monitored the location of eGFP-uS11 in various Krr1 mutant strains. As observed in cells complemented with Krr1, the viable Krr1ΔC50 and Krr1* mutants displayed cytoplasmic localisation of eGFP-uS11, indicating nuclear incorporation of the r-protein and proper export of pre-ribosomes into the cytoplasm (Fig. 7c, panels 1-3). By contrast, the Krr1*ΔC50 mutation induced a strong nucleolar/nuclear accumulation of eGFP-uS11, pointing to an early assembly defect (Fig. 7c, panel 4). Interestingly, the viable Krr1ΔC50 mutant, but not the Krr1* mutant, specifically showed cytoplasmic accumulation of ITS1 RNA, as judged by FISH experiments (Fig. 7d). This phenotype was not observed in the Krr1*ΔC50 expressing strain, suggesting that disruption of the PRP in the Krr1ΔC50 mutant induces an earlier assembly block that precedes, and therefore masks, cytoplasmic ITS1 RNA accumulation observed in Krr1ΔC50 cells (Fig. 7d). A cryo-EM structure of an early pre-A1 state of the 90S pre-ribosome showed that Krr1^PRP^ also provides a binding platform for the r-protein uS15 (yeast Rps13)^56^ (Fig. 7e). We propose that, through a dynamic exchange of partners, Krr1^CTT^ coordinates multiple events during 90S pre-ribosome assembly.

## Discussion

In eukaryotes nascent non-native r-proteins must be shielded from aggregation, imported into the nucleus through nuclear pores, and maintained in assembly-competent state prior to their incorporation into pre-ribosomes. Although multiple importins and dedicated chaperones perform these steps, how these delivery factors interface with the assembly machinery to precisely deposit r-proteins remains poorly understood. Capturing and defining these connections is challenging as the underlying interactions are transient and not stably retained on affinity purified pre-ribosomes. Fap7, a nuclear localised isoaspartylase essential for the conformational maturation of uS11, is one such factor that does not co-enrich with pre-ribosomes^21^. However, its depletion induces cytoplasmic 20S pre-rRNA processing defects, creating an apparent mismatch between its inferred site of action and its mutant phenotype^21,57,58^.

Here, we place Fap7 within a nucleolar assembly-factor network that drives 90S pre-ribosome formation. TurboID-based proximity labelling revealed that Fap7 resides in an environment that enriched not only uS11 but also Krr1, Kri1, Imp3, and Faf1 - all factors that dock on the 90S pre-ribosome. Within this network, we show that Krr1 is the principal assembly factor that binds to Fap7. Fap7 recognizes a conserved PRP within Krr1^CTT^ through a cavity formed within the C-terminal region of Fap7. Our crystal structure of CtFap7-PRP defines this interaction at high-resolution and provides the basis for its conservation. Comparison with archaeal homologs of Fap7 indicates that this interaction platform is a eukaryotic innovation. The archaeal Fap7 family retains uS11-binding activity but lacks the C-terminal architecture required for Krr1 engagement and accordingly fails to complement the lethality of Fap7 depletion in yeast. By contrast, TbFap7, which contains a diverged Krr1-binding pocket and exhibits a weakened Krr1 interaction, rescues viability to nearly wild-type levels when overexpressed. We suggest that the ancestral chaperoning and isoaspatylase activity toward uS11 were retained, while the acquisition of a Krr1-binding surface embedded Fap7 into a conserved nucleolar assembly-factor network. The presence of the PRP and the Fap7’s C-terminal binding pocket across eukaryotes, together with their absence in archaea, is consistent with their co-evolution.

Functional expansion enabled the ancestral Fap7 to also function as a licensing factor within a hierarchically assembled module. Fap7 engages with the Krr1^CTT^ and an additional surface on the KH2 domain. In the absence of uS11, this arrangement gates the access of Kri1 to Krr1 due to steric clashes with the bound Fap7. uS11 incorporation relieves this barrier, i.e. protection of the KH2 region is lost to permit Kri1 interaction. These studies support a sequential pathway in which a pre-assembled Krr1:Fap7 complex captures uS11 and then transitions into a Kri1-binding competent state. In this framework, Fap7 couples r-protein cargo loading to subsequent assembly-factor recruitment, establishing an early checkpoint that imposes directionality to the assembly process. Our TurboID data indicate that the environment of Fap7 encompasses the Krr1-proximal assembly factors Imp3 and Faf1 (Fig. 2a), suggesting that these dynamic events occur in the vicinity of the 90S pre-ribosome prior to the installation of snR30 snoRNP satellite / uS15 incorporation.

Our functional studies clarify the role of Fap7 during 40S subunit assembly. We propose that isoAsp modification by Fap7 alters the backbone geometry of uS11^CTT^ ^22^. The equivalent aspartate in yeast mitochondrial uS11m is occupied by glycine, an alternative solution to achieve the required local backbone geometry. Glycine uniquely accesses a broader region of the Ramachandran space^59,60^, raising the possibility that a glycine substitution structurally mimics an isoAsp. Consistent with this model, substitution of aspartate to glycine within uS11 bypassed the requirement for Fap7 and restored 20S pre-rRNA processing defects observed upon Fap7 depletion. Thus, isoaspartylation of Asp124 in uS11 or glycine at the equivalent position in yeast uS11m represent distinct mechanisms to impose a similar uS11^CTT^ conformation necessary for correct 40S subunit assembly. The dominant nature of the suppression suggests that final maturation does not depend on the continued presence of Fap7 itself on the 40S pre-ribosome, but on establishment of the Fap7-induced uS11^CTT^ conformation, which serves as a late cytoplasmic checkpoint.

Whereas uS11 depletion blocks early pre-rRNA processing^57,58^, Fap7 depletion causes cytoplasmic accumulation of 20S pre-rRNA, consistent with impaired cleavage at site D. We propose that without Asp124 modification, the local backbone of uS11^CTT^ fails to adopt the conformation needed to correctly dock rRNA helices h23, h24, and h45 within the platform. This structural defect remains latent during nucleolar assembly but becomes exposed only at later quality control steps in the cytoplasm, when platform architecture is evaluated during 20S pre-rRNA processing steps. Notably, several substitutions within uS11^CTT^ similarly lead to 20S pre-rRNA accumulation phenocopying Fap7 depletion^61–63^. As this region is functionally coupled to the pathway that permits Nob1 access^64^, we suggest that defective docking of the rRNA helices onto uS11^CTT^ may underlie the observed block in cleavage at site D.

Our findings implicate Krr1^CTT^ as a multifunctional coordinator of 90S pre-ribosome assembly. In addition to recruiting Fap7, the Krr1^PRP^ provides a binding platform for the r-protein uS15 and contributes to the release of the snR30 snoRNP^37,41,56^. This versatility may explain why the Krr1ΔC70 truncation mutant exhibits early nucleolar defects that obscure later consequences of impaired Fap7 recruitment^37^. We suggest that Krr1^CTT^ integrates multiple assembly events, with Fap7-mediated uS11 conformational maturation representing one task.

In conclusion, this work illustrates how an archaeal enzyme was embedded into the complex eukaryotic pathway: its ancestral activity was retained, while a new interaction surface integrated it into a eukaryotic assembly-factor network. The discovery of a Fap7-centred module provides a framework for understanding how non-native nascent r-proteins, in general, may be safely guided to their cognate rRNA-binding sites, with their incorporation being tightly coordinated by specific assembly factors and with ongoing maturation events. We suggest that formation of transient modules introduces checkpoints that ensure ordered co-delivery of nascent r-proteins and assembly factors to developing pre-ribosomes, thereby safeguarding the fidelity of ribosome production. Such checkpoints echo regulatory mechanisms observed in other archaeal-eukaryotic pathways. In the ESCRT pathway, the ancestral membrane-remodelling ATPase Vps4 became integrated through eukaryotic specific ESCRT-III adaptors into a regulated, ATP-dependent cycle^65–67^. The cytoskeletal systems follow a related logic: actin-family proteins, derived from archaeal actin-like ancestors, were incorporated into eukaryotic regulatory networks via actin-binding proteins, making polymerization dependent on spatial and signalling cues^68,69^. Thus, the Fap7-centred module reflects a recurrent evolutionary strategy in which ancient enzymatic activities are embedded into interaction-dependent circuits to impose order, timing, and fidelity of complex eukaryotic pathways.

## Acknowledgements

We thank Matthias Peter for sharing yeast strain collections, R. Pillai and the Panse laboratory for discussions. OV & VGP thank the Proteomics Core Facility, A. Hainard and R. Visentin from the Faculty of Medicine, University of Geneva for HDX-MS data acquisition. SF & DK thank Michael Stumpe of the Proteomics Unit of the Metabolomics and Proteomics Platform (MAPP), University of Fribourg for his support and assistance. We thank Andrew McCarthy, EMBL Grenoble, and the EMBL-ESRF beamline staff at ESRF, Grenoble, for crystal mounting and data collection.

## Author contributions

Experimental design: AGG, SF, OV, YV, RS, MAR, LZ, CH, MO-O, AS, DK & VGP. Experiment execution: AGG, SF, OV, FA, TvA, YV, RS, MAR, LZ, CH, MO-O & DK. Data analysis: AGG, SF, OV, FA, TvA, RS, MAR, LZ, CH, MO-O, DP-C, DK & VGP. Supervision: MO-O, MS, PB, DK, VGP. Writing-original draft: AGG & VGP. Writing-review and editing: AGG, SF, OV, FA, TvA, MO-O, DP-C, MS, PB, AS, DK & VGP

## Methods

### Yeast strains and plasmids

All *Saccharomyces cerevisiae* strains used in this study are listed in Supplementary Table 2. Genomic disruptions, promoter switches, and C-terminal tagging at genomic loci were performed using standard yeast molecular biology^70^. All plasmids used in this study are listed in Supplementary Table 3. All recombinant DNA work was performed using *Escherichia coli* TOP10 cells. Gene mutations were generated with QuickChange site-directed mutagenesis kit (Agilent Technologies, Santa Clara, CA, USA). All cloned DNA fragments and mutagenized plasmids were verified by sequencing.

### Yeast two-hybrid interaction assays

Full-length and/or fragments of Krr1, yeast Fap7, TbFap7, and Faf1 were fused to either a LexA DNA-binding domain (BD; pLexA *TRP1* marker) or a Gal4 activation domain (AD; pACT2.2 *LEU2* marker) and co-transformed into the NMY32 reporter strain, containing the *HIS3* and *ADE2* reporter genes. Transformants were grown on selective SD plates, lacking leucine and tryptophan (SD-Leu-Trp). Cells were spotted from 10-fold serial dilutions onto the corresponding media (SD-Leu-Trp, SD-His or SD-Ade) and incubated for 3 days at 30°C. The strength of the interaction on SD-His plates was assessed using increasing concentrations of 3-amino-1,2,4-triazole (3-AT; Merck), added to the medium. 3-AT competitively inhibits the product of imidazole glycerol-phosphate dehydratase, thereby increasing the selective pressure on *HIS3* activation. SV40 large T antigen (LargeT) and human lamin C (LaminC) were used as negative controls.

### TurboID-based proximity labelling assay

Plasmids expressing N- or C-terminally TurboID-tagged Fap7 and the appropriate control proteins under the control of the copper-inducible *CUP1* promoter were transformed into the wild-type yeast strain YDK11-5A^71^. The TurboID-based proximity labelling experiment, sample processing, and data acquisition by LC-MS/MS were performed as previously described^36^. LC-MS/MS measurements were performed on a Q Exactive HF-X (Thermo Scientific) coupled to an EASY-nLC 1200 nanoflow-HPLC (Thermo Scientific). The MS raw data files were analysed with the MaxQuant software package version 1.6.2.10^72^ for peak detection, generation of peak lists of mass-error-corrected peptides, and database searches as previously described^36^. For quantification, missing iBAQ (intensity-based absolute quantification) values in the two control purifications from cells expressing either GFP-TurboID or NLS-GFP-TurboID (controls for the experiment with C-terminally TurboID-tagged Fap7) or TurboID-GFP or NLS-TurboID-GFP (controls for the experiment with N-terminally TurboID-tagged Fap7) were imputed in Perseus^73^. For normalization of intensities in each independent purification, iBAQ values were divided by the median iBAQ value, derived from all nonzero values, of the respective purification. To calculate the enrichment of a given protein compared to its average abundance in the two control purifications, the normalized iBAQ values were log_2_-transformed and those of the control purifications were subtracted from the ones of the Fap7 bait purification. For graphical presentation, the normalized iBAQ value (log_10_ scale) of each protein detected in the Fap7 bait purification was plotted against its relative abundance (log_2_-transformed enrichment) compared to the control purifications. The datasets have been deposited to the ProteomeXchange Consortium via the PRIDE^74^ repository with the dataset identifier (PXD078992). Project accession: PXD078992; Token: jyukJLn9ZW5q

### Recombinant protein expression and purification

Recombinant proteins were expressed in *E. coli* BL21(DE3) cells at 20°C overnight, by 0.5mM IPTG (Panreac Applichem) (added at OD_600_ 0.6-1). Cells were resuspended in buffer containing 300mM NaCl, 50mM HEPES, pH 7.5 (unless stated otherwise) and lysed either with a microfluidizer (version M-110P, Microfluidics), three times at 20,000psi, or by sonication (Sonic Ruptor 4000, Omni International) for three cycles at 75% of the maximum output for 3 min/cycle. Lysates were cleared by centrifugation at 38,750 ×g for 10min, and the supernatant was collected. Purification proceeded using either Nickel-NTA beads (Ni^2+^-NTA; Qiagen) or Glutathione Sepharose 4 Fast Flow beads (GSH; Cytiva), according to protein requirements. Ni-NTA beads or GSH beads were added to the supernatant and incubated for 1h at 4 °C. The samples were then loaded into a plastic column fitted with a polyethylene frit (Merck). Ni^2+^-NTA beads were washed three times with buffer supplemented with 30 mM imidazole (Thermo Fisher), while GSH beads were washed with lysis buffer supplemented with 0.1% Tween-20 (Thermo Fisher). The bound protein was eluted by 300–500 mM imidazole (histidine-tagged proteins) or 50mM glutathione (GST-tagged proteins; PanReac AppliChem) and dialysed overnight at 4 °C. The protein was concentrated using spin filter tubes (Amicon, Merck) and stored at −80°C with 10% glycerol. Whenever Krr1 full-length samples were used, bound RNA was removed by cation exchange chromatography. The eluate was dialyzed overnight at 4°C against 1l of DIA-Buffer (400 mM NaCl, 50 mM HEPES, pH 7.5, 5% glycerol, 3 mM DTT). The protein was then mixed with a lower salt buffer (300mM NaCl, 25mM HEPES, pH 7.5) and loaded onto a HiTrap SP HP cation exchange chromatography column (Cytiva) using a peristaltic pump. A salt gradient from 300 mM to 1M NaCl was applied using an ÄKTA Pure system (Cytiva). RNA-free Krr1 typically eluted at a conductivity of 45-5mS/cm. When removal of the His-tag was required, TEV protease digestion was performed overnight, followed by incubation with Ni^2+^-NTA beads to bind the cleaved tag and TEV protease and let the protein run through. The protein was concentrated using spin filter tubes (Amicon, Merck) and stored at −80°C with 10% glycerol.

### Biochemical interaction studies

Binding assays were performed by mixing 50μl of bead slurry affinity beads (either Ni^2+^-NTA or GSH beads) with 100-500pmol of purified bait proteins (either His_6_- or GST-tagged) and equimolar amounts of untagged prey proteins in a 1.5 mL tube. 500μl of buffer was added (either PBS with 0.1% Tween-20; HBS with 0.1% Tween-20; or HEPES pH 7.5, 300mM NaCl, 0.1% Tween-20 supplemented with 30mM imidazole when working with Ni^2+^-NTA beads). The proteins were incubated at 4 °C, for 30min, rotating. The beads were pelleted by centrifugation at 870 ×g and the supernatant was discarded. The beads were then washed three times with 500μl of buffer and centrifuged in the previous conditions. Proteins were eluted with 25μl of 2×LDS buffer (Novex), heated at 95 °C for 10 min, separated by SDS-PAGE and visualized by Coomassie Brilliant Blue staining.

### Hydrogen-Deuterium Exchange mass spectrometry (HDX-MS)

HDX-MS experiments were performed at the Protein Biochemistry Platform of the University of Geneva following established protocols with minimal modifications^52^. Protein concentrations and their ratios, details of reaction conditions and all the data are presented in Supplementary Data 1 & 2. HDX reactions were performed in 50µl volumes with a final protein or protein-complex concentration of 2 or 4.2μM. Briefly, deuterium exchange reaction was initiated by adding 40µl of D_2_O exchange buffer to the protein sample. Reactions were carried-out on ice for three incubation times (3 s, 30s, 300s) and terminated by the sequential addition of 20µl of ice-cold quench buffer 1 (4M Gdn-HCl, 1M NaCl, 100 mM NaH_2_PO_4_, pH 2.4, 1% formic Acid, FA). Samples were immediately frozen in liquid nitrogen and stored at −80°C for up to two weeks. All experiments were repeated in triplicate. To quantify deuterium uptake into the protein, samples were thawed and injected in a UPLC system immersed in ice with 0.1% FA as liquid phase. The protein was digested via two immobilized pepsin columns (Thermo Fisher Scientific), and peptides were collected onto a VanGuard precolumn trap (Waters). The trap was subsequently eluted, and peptides separated with a C18, 300Å, 1.7μm particle size Fortis Bio 100 × 2.1mm column over a gradient of 8-30% buffer C over 20min at 150ml/min (buffer B: 0.1% FA; buffer C: 100% acetonitrile). Mass spectra were acquired on an Orbitrap Velos Pro (Thermo Fisher Scientific), for ions from 400 to 2200 m/z using an electrospray ionization source operated at 300 °C, 5 kV of ion spray voltage. Peptides were identified by data-dependent acquisition of a non-deuterated sample after MS/MS and data were analysed by Mascot. All peptides analysed are shown in Source Data Table 1-2. Deuterium incorporation levels were quantified using HD examiner software (Sierra Analytics), and the quality of every peptide was checked manually. Results are presented as percentage of maximal deuteration compared to theoretical maximal deuteration. Changes in deuteration level between the two states were considered significant if >9% and >0.7 Da and *p* < 0.05 (unpaired t-test). The dataset has been deposited into the Yareta Database hosted by University of Geneva (https://yareta.unige.ch/home) can be accessed via: https://doi.org/10.26037/yareta:yxlxksvpzvg45bd2fxivq4azey (DOI: 10/hb7bs2).

### CtFap7-PRP purification for structural studies

*Ct*Fap7 (from which amino acids 2-6 were removed to facilitate crystallization) and a *Ct*Fap7 variant in which the consensus sequence of the PRP was fused to its C-terminus via a GSG-linker (*Ct*Fap7-PRP) were purified and crystallized. BL21(DE3) *E. coli* cells expressing *Ct*Fap7-PRP were grown in 4L of LB medium. The cells were harvested and lysed using lysis buffer (50mM HEPES pH 7.5, 300mM NaCl) and two spatula tips of lysozyme (Thermo Fisher). The lysate was sonicated for three cycles of 3min each at 70% power (Sonic Ruptor 4000, Omni International). The lysate was loaded onto 4ml of Ni^2+^-NTA (Qiagen) resin. The beads were washed with lysis buffer, followed by wash buffer (50mM HEPES pH 7.5, 300mM NaCl, 50mM imidazole). The protein was eluted with 2 × 10ml of elution buffer containing 500 mM imidazole. The His_6_-tag was removed by the addition of 200µg TEV protease and dialyzed against 1 litre of dialysis buffer (HBS + 3mM DTT). The supernatant was loaded onto 5ml of Ni^2+^-NTA resin (Qiagen) and the flow-through was collected. The protein was concentrated to ∼2ml using centrifugal concentrators (Amicon, Sigma-Aldrich) and subjected to size-exclusion chromatography (SEC) using an ÄKTA system and a Superdex 200 Increase 10/300 GL column (Cytiva), equilibrated with buffer containing 400mM NaCl, 25mM HEPES pH 7.5, 3mM DTT, and 5mM MgCl₂. Fractions containing the protein of interest were pooled and concentrated to ∼15mg/ml using an Amicon concentrator (Sigma-Aldrich). AMP-PNP (Sigma-Aldrich) was added to a final concentration of 10mM.

### Crystallization and structure determination

Crystallization screening was performed at the Protein Crystallization Centre, UZH. For *Ct*Fap7-PRP, 0.2µl of ∼15 mg/ml protein was mixed with 0.2µl of reservoir solution (75 µl well size) and incubated at 20°C and 4°C using the sitting-drop vapor diffusion method. Crystals formed within two days at 20°C in two conditions: (1) 0.2M MgCl₂, 0.1M TRIS pH 8.5, 30% (w/v) PEG 4K; (2) 0.2M sodium acetate, 0.1 M TRIS pH 8.5, 30% (w/v) PEG 4K. The crystals were cryoprotected with reservoir solution + 25% (v/v) glycerol and flash-frozen in liquid nitrogen. Diffraction data for *Ct*Fap7-PRP were collected at the European Synchrotron Radiation Facility (ESRF) at 100K (EMBL, Grenoble, France) and processed in space group P2_1_. Two copies of the *Ct*Fap7-PRP complex were present in the asymmetric unit. Phases were determined by molecular replacement using the Phaser module^75^ of the Phenix package^76^, with *Ct*Fap7-PRP models generated by AlphaFold2-based ColabFold as initial search models^49^. Model building was performed manually in Coot^77^, and refinement was carried out using the Phenix refine module^78^.

### Protein structure modelling and sequence analysis

Single- and multi-chain protein structure predictions were generated using AlphaFold2 v1.5.5 via the Google Colab platform^49,79^. For each prediction, model confidence was assessed using the predicted Local-Distance Difference Test (pLDDT) scores, Predicted Aligned Errors (PAE), and pTM/ipTM scores^49,80^. The highest-confidence models were visualized using PyMOL v2.5.5 or UCSF ChimeraX 1.8^81^ software. Protein sequences were retrieved from the SGD^82^ and UniProt^83^ databases. Sequence alignments were generated in Jalview v2.11.4.1 using its integrated ClustalO algorithm^84,85^.

### Molecular dynamics simulations

Molecular dynamics simulations were performed for the two peptide sequences PSDST and PSGST. Initial peptide models were generated using the AlphaFold 3 Server^50^. System preparation and simulations were carried out with GROMACS 2026.1^86^ using the AMBER99SB-ILDN force field^87^ and the TIP3P water model. Each peptide was placed in a rhombic dodecahedron box with a minimum solute-box distance of 1.4nm, solvated, and neutralized with 0.15M NaCl. After energy minimization, each system was equilibrated for 100 ps in the NVT ensemble at 310 K with heavy-atom position restraints of 1000kJmol⁻¹ nm⁻². This was followed by sequential 100ps NPT equilibration steps at 310K and 1bar, during which restraints were reduced stepwise from 1000 to 100 and 10 kJ mol⁻¹ nm⁻², followed by a final 100 ps unrestrained NPT equilibration. Production simulations were performed for 50 ns in the NPT ensemble at 310K and 1bar using a 2fs time step. Bond lengths were constrained using LINCS, short-range non-bonded interactions were truncated at 1.0 nm, and long-range electrostatics were treated with the particle mesh Ewald method. Coordinates were saved every 0.5ns. Trajectory analysis was performed on 5000 extracted frames. Backbone dihedral angles, φ and ψ, were calculated for the central Asp residue in -PSDST- or the Gly residue in -PSGST-. The φ/ψ distributions were visualized as Ramachandran maps using Matplotlib^88^.

### Monitoring 20S pre-rRNA by fluorescence in situ hybridization (FISH)

FISH experiments were performed as described previously^89^. Yeast cultures (50ml) were grown to OD_600_ 0.7 at 25 / 30°C. Formaldehyde was added to a final concentration of 4% (v/v), and incubation continued for 15min with gentle shaking. Cells were harvested by centrifugation at 3,300rpm for 1min (Beckman JA25.5 rotor) and resuspended in 50ml of fixation buffer (0.1M potassium phosphate buffer, pH 6.4, 4% formaldehyde). The suspension was incubated at room temperature for 1h, on a rotating wheel. Cells were pelleted by centrifugation at 3,300 rpm for 1min, resuspended in 1ml of 0.1M potassium phosphate buffer, pH 6.4, and transferred to 1.5ml tubes. After centrifugation (3,300rpm, 1min), cells were washed once with 1 mL of potassium phosphate buffer (pH 6.4) and twice with 1ml of wash buffer (0.1M potassium phosphate, 1.2M sorbitol, pH 6.4). Pellets were stored overnight at 4°C. The next day, cell pellets were resuspended in 1 ml of wash buffer containing 500μg zymolyase 100T, 10mM DTT, and 10mM ribonucleoside vanadyl complex (RVC), then incubated at 30°C on a rotating platform for 30min to induce spheroplast formation. Spheroplasts were collected by centrifugation at 3,000rpm for 3min, resuspended in 1ml of wash buffer, centrifuged again under the same conditions, and resuspended in a volume of wash buffer twice that of the pellet. Slides (Thermo Scientific, Menzel-Gläser, Diagnostic slides, 8-well, 6mm) were coated with poly-L-lysine solution by immersion and incubated on a shaking platform for 5min before removing most of the solution. Slides were left to dry overnight at room temperature. 20μL of washed cells were applied to each well and incubated for 5min at room temperature. Non-adherent cells were removed by aspirating excess liquid. The slides were washed with 2× SSC (0.3M NaCl, 0.03M trisodium citrate pH 7.4), followed by a second wash with 2× SSC and a 10-min incubation. 12μl of pre-hybridization buffer (PBS containing 50% formamide, 10% dextran sulfate, 4× SSC, 0.02% polyvinylpyrrolidone, 0.02% bovine serum albumin, 0.02% Ficoll-400, 125μg/mL *E. coli* tRNA, 500μg/ml herring sperm DNA, 10mM RVC, and 1mM DTT) was added to each well. Slides were incubated overnight at 37°C in a humid chamber. Hybridization was performed by incubating 5pmol of Cy3-labeled DNA probe complementary to the ITS1 (5’-Cy3-ATGCTCTTGCCAAAACAAAAAAATCCATTTTCAAAATTATTAAATTTCT-3’) to each well, avoiding light exposure. The slides were incubated overnight at 37°C in a humid chamber. Post-hybridization washes were carried out with 2× SSC and 1× SSC, each with 5-min incubation on a shaking platform. DNA was stained with DAPI (10μg/mL in 1× SSC) for 5 min, followed by a wash with 1× SSC and incubation in 0.5× SSC for 30min. Slides were dried for 2 h before mounting with Mowiol mounting medium. Imaging was performed using a THUNDER Imager 3D Assay (Leica) equipped with an HCX PL-APO FLUOTAR ×63/0.60 NA oil immersion objective.

### Fluorescence microscopy

Yeast cells carrying the indicated GFP reporters were grown in the appropriate selective medium prior to imaging. Unless otherwise stated, transformants were plated and re-streaked on selective synthetic medium, grown overnight at 25 / 30°C with shaking at 150 rpm, diluted to OD_600_ = 0.2, and grown to mid-log phase. For P_GAL_-KRR1 strains co-expressing eGFP-uS11 with KRR1, KRR1ΔC50, KRR1* or KRR1*ΔC50, cells were selected on SG-Ura, re-streaked and grown in SD-Ura before imaging. P_GAL_-KRR1 cells co-expressing eGFP-uS11 and KRR1*ΔC50 were selected on SG-Ura and grown in SD-Ura for approximately 36h at 30°C before microscopy. Wild-type cells expressing eGFP-uS11, were grown in SD-Ura, whereas cells expressing yeast Fap7-GFP or aFap7-GFP were grown in SD-His or SD-Leu, respectively. Cells were harvested by centrifugation for 3min at 2300 rpm, resuspended in water, and 4µl of cell suspension was mounted in the centre of an ibidi dish and immobilized with 4% agarose pad and covered with a coverslip (18×18mm No. 1, VWR). Cells were visualized using a Leica DMi8 Thunder Imager 3D microscope (Leica, Germany) equipped with an HC PL APO 63x/1.40-0.60 oil immersion objective (Leica, Germany). Images were acquired with a Leica DFC9000 GT digital camera (Leica, UK) and processed using ImageJ software (version 1.54p; NIH and LOCI, USA).

### Tandem-affinity purification

A conditional P_GAL_-uS11 mutant expressing C-terminally FTpA-tagged (3xFLAG-TEV-ProteinA) Krr1 were grown in 2-4L of YPG (+uS11) and YPD (-uS11) media. Cells were harvested at 5,000rpm for 20min at 4°C. The cell pellet was resuspended in 25ml of lysis buffer (50mM TRIS pH 7.5, 75mM NaCl, 1.5mM MgCl_2_, 0.15% (v/v) Igepal CA-630) containing 1 mM DTT, 1.5 mM PMSF and ½ tablet of complete EDTA-free protease inhibitor cocktail (Roche). Cells were lysed at 500rpm for 20min at 4°C with glass beads (400-600 mm diameter) in a Pulverisette 6 planetary mill (Fritsch). Lysates were clarified by centrifugation at 4°C for 20 and 30min at 5,000rpm and 15,000rpm, respectively. Lysates were then incubated with 300 µl of preequilibrated IgG-Sepharose beads (Cytiva) for 1.5h at 4°C. Post IgG incubation, beads were collected in a 10ml chromatography column (Bio-Rad Laboratories) and washed twice with 5 ml of lysis buffer containing 0.5 mM DTT. The bound pre-40S particles were eluted from the IgG beads by incubating in 1ml of lysis buffer containing 0.5mM DTT with TEV (tobacco etch virus) protease at 4°C overnight. TEV eluates were collected in a 10ml chromatography column, and IgG beads were washed with 200µl of lysis buffer twice and collected in the same column (total of 1.4ml). 100 µl of preequilibrated FLAG beads (Cytiva) were added to the TEV eluates to allow binding of pre-ribosomes, supplemented with 0.5mM DTT. This step was performed on a rotating wheel, at 4°C for 2h. Post incubation, beads were washed three times with 1 ml of lysis buffer and elution was performed twice with 150 µl of elution buffer containing 50 mM TRIS pH 7.5, 150mM NaCl, and 100mg/ml of 3 × FLAG^TM^ peptide (Merck) at room temperature for 30min. The eluates were TCA precipitated and washed twice with ice-cold acetone. The final pellet was dissolved in 1 × LDS sample buffer (Invitrogen) and samples were analysed with silver staining and western blotting after separation on NuPAGE 4-12% Bis-Tris gradient gels (Invitrogen) and SurePAGE Bis-Tris 4-20% gradient gels (Genscript, EUA), respectively.

### Mass spectrometry

Protein pellets were resuspended in 500µl of 100mM ammonium bicarbonate (pH 7.5), then reduced by addition of 5mM dithiothreitol for 60 min at 37°C and alkylated with 12mM iodoacetamide for 60 min in the dark. Overnight digestion was performed with trypsin (Promega, sequencing-grade) at a 1:100 proteinase: protein (w/w) ratio at 37 °C with gentle agitation. Resultant peptides were acidified by the addition of 3% (v/v) formic acid and then desalted using BioPureSPN C18 plates (HNS S18-V-L, Macro-96-Well, NEST Group). To remove the FLAG peptide, plates were washed three times with 12% (v/v) acetonitrile / 0.1% (v/v) formic acid prior to elution with 300µl of 50% (v/v) acetonitrile / 0.1% (v/v) formic acid. Peptides were then dried in a SpeedVac before resuspension in 30µl of 0.1% (v/v) formic acid for analysis by liquid chromatography-mass spectrometry.

Peptides were analysed using data-dependent acquisition (DDA) on an Orbitrap Exploris 480 Mass Spectrometer (Thermo Fisher Scientific) coupled to a Vanquish Neo UHPLC System (Thermo Fisher Scientific). Liquid chromatography was performed with a 60-min gradient (flow rate = 300nl/min) on a 40cm × 0.75mm column (CoAnn ICT36007515F-50-5) packed in-house with C18 beads (Dr. Maisch, Reprosil-Pur 120). Precursor scans were acquired in the Orbitrap with the following settings: resolution = 120,000; scan range = 350 – 1500*m*/*z*; normalised AGC target = 200%; maximum injection time = 100ms. Precursor ions were subsequently selected for fragmentation by higher-energy collisional dissociation (HCD) using a normalised collision energy of 30%. Fragment ions were acquired in the Orbitrap with the following settings: isolation window = 1.4*m/z*; normalised AGC target = 200%, maximum injection time = 54ms. Mass spectrometry raw files were analysed using the MSFragger^90^ and IonQuant^91^ plug-ins in FragPipe v19.0. Data were searched against the *Saccharomyces cerevisiae* proteome (UniProt ID: UP000002311; 6177 entries), supplemented with common contaminants and reverse decoys for false-discovery rate (FDR) estimation. The “Default” workflow was used with carbamidomethylation of cysteine (+57.02146 Da) set as a fixed modification and oxidation of methionine (+15.9949 Da) set as a variable modification. Label-free quantification of MS1 peptide intensities was performed using IonQuant with match-between-runs (MBR) and MaxLFQ enabled, followed by normalization across runs using unique and razor peptides. Output files were further analysed and visualized using FragPipe-Analyst^92^. The dataset has been deposited to the ProteomeXchange Consortium via the PRIDE partner repository^74^ with the dataset identifier PXD079433. Project accession: PXD079433 Token: Y36o5nrO3w6z.

## Figures

Figures of molecular structures were created using PyMOL v2.2.0 or ChimeraX 1.8 software. Sequences were searched in the SGD, UniProt, and BLAST databases. Sequence alignments were generated using MAFFT (v7) and JalView (2.11.1.4).

## Statistics and reproducibility

All data are representative results from at least three independent experiments unless otherwise specified. No statistical method was used to predetermine the sample size. The experiments were not randomized. The investigators were not blinded to allocation during the experiments and outcome assessment. Statistical analysis was performed using Prism (version 10.3.0; GraphPad Software Inc., La Jolla, CA, USA) or Microsoft Excel (Microsoft Office, version 16; Microsoft Corporation, Redmond, WA, USA).

## Data availability

All data are presented in the main text and Figures or Supplementary information can be requested from the corresponding author. The TurboID datasets generated in this study have been deposited in the ProteomeXchange Consortium via the PRIDE partner repository^74^ under accession code PXD078992. Proteomes of Krr1-FTpA in wild-type and uS11 depletion mutant have been deposited in the ProteomeXchange Consortium via the PRIDE partner repository ^74^ under accession code PXD079433. HDX-MS data have been deposited in the Yareta Database hosted by the University of Geneva (https://yareta.unige.ch/home) can be accessed via the DOI:10/hb7bs2. The peptides used for HDX-MS analyses are provided as Supplementary Data files 1 and 2. Coordinates for the crystal structures of Fap7 and the Fap7 complex have been deposited in the Protein Data Bank (PDB) under accession code 31GO (Deposition code ID D_129215714). All strains and plasmids used in this study are listed in Supplementary Tables 2 and 3. Detailed protocols, plasmid maps and oligonucleotide-based cloning/mutagenesis strategies are available upon reasonable request. Supplementary Source data are provided with this paper.

