## Supplemental Information for "A nucleolar assembly module integrates an ancestral isoaspartylase to safeguard ribosome biogenesis"

**Table of Contents**

1. Supplementary Tables 1 to 3 (p. 2-4)
2. Supplementary Figures 1 to 2 (p. 5-6)
3. Uncropped SDS-PAGE gels (p.7)
4. Supplementary References (p.8)

**Supplementary Table 1. Crystal structure information table.**

| | *CtFap7 complexed with Krr1-PRP* |
|---|---|
| **Data collection** | |
| **Space group** | P 1 21 1 |
| **Cell dimensions** | |
| **a, b, c (Å)** | 38.655 34.873 146.737 |
| **α, β, γ (°)** | 90 93.347 90 |
| **Wavelength (Å)** | 1 |
| **Resolution range (Å)** | 146.5 - 1.78 (1.844 - 1.78) |
| **Total reflections** | 70255 (5997) |
| **Unique reflections** | 36121 (3097) |
| **Multiplicity** | 1.9 (1.9) |
| **Completeness (%)** | 94.71 (81.51) |
| **Mean I/sigma(I)** | 5.69 (1.29) |
| **Wilson B-factor** | 27.35 |
| **R-merge** | 0.04301 (0.4206) |
| **R-meas** | 0.06083 (0.5948) |
| **R-pim** | 0.04301 (0.4206) |
| **CC1/2** | 0.998 (0.807) |
| **CC*** | 0.999 (0.945) |
| **Refinement** | |
| **Reflections used in refinement** | 36063 (3091) |
| **Reflections used for R-free** | 1741 (139) |
| **R-work** | 0.2167 (0.3874) |
| **R-free** | 0.2357 (0.4134) |
| **CC(work)** | 0.949 (0.806) |
| **CC(free)** | 0.936 (0.723) |
| **Number of non-hydrogen atoms** | 3240 |
| **Macromolecules** | 2988 |
| **Ligand** | 90 |
| **Solvent** | 188 |
| ***R.M.S. deviations*** | |
| **Bond lengths (Å)** | 0.014 |
| **Bond angles (°)** | 1.73 |
| **Ramachandran favored (%)** | 97.83 |
| **Ramachandran allowed (%)** | 2.17 |
| **Ramachandran outliers (%)** | 0.00 |
| **Rotamer outliers (%)** | 1.51 |
| **Clashscore** | 7.03 |
| **Average B-factor** | 39.07 |
| **Macromolecules** | 39.12 |
| **Ligand** | 33.04 |
| **Solvent** | 40.34 |

**Supplementary Table 2.** ***Saccharomyces cerevisiae*** **strains used in this study.**

| Name | Genotype | Origin |
|---|---|---|
| BY4741 | *MATa; his3-1; leu2-0; met15-0; ura3-0 TRP+* | Euroscarf |
| NMY32 | *MATa trp1 leu2 (lexAop)8-ADE2 LYS2::(lexAop)4-HIS3 URA3::(lexAop)8-lacZ GAL4* | Dualsystems Biotech AG |
| $P_{GAL1}$-Krr1 | BY4741, $P_{GAL1}$-*KRR1*::NAT | This study |
| $P_{GAL1}$-Fap7 | BY4741, $P_{GAL1}$-*FAP7*::NAT | Peña et al. (2016)[1] |
| $P_{GAL1}$-uS11 | *Mat alpha his3-200 trp1-101 leu2-2 ura3-167 rps14a::kanMX6 rps14b::LEU2 + pGAL-RPS14A URA3* | Jakovljevic et al. (2004)[2] |
| yrb2Δ | *BY4741, YRB2::kanMX6* | Oborská-Oplová et al. (2022)[3] |

**Supplementary Table 3. Plasmids used in this study.**

| Name | Relevant markers | Source |
|---|---|---|
| pLexA Faf1 | *LexA-FAF1 2µ TRP1* $AMP^R$ | This study |
| pLexA Fap7 | *LexA-FAP7 2µ TRP1* $AMP^R$ | This study |
| pLexA SiFap7 | *LexA-SiFAP7 2µ TRP1* $AMP^R$ | GenScript |
| pLexA TbFap7 | *LexA-TbFAP7 2µ TRP1* $AMP^R$ | Twist Bioscience |
| pLexA LaminC | *LexA-LAMINC 2µ TRP1* $AMP^R$ | Dual Systems |
| pAct2.2 Krr1 CTT/193-316 | *GAL4 AD-KRR1* 193-316 *2µ LEU2* $AMP^R$ | This study |
| pAct2.2 Krr1 193-266 | *GAL4 AD-KRR1* 193-316 267stop *2µ LEU2* $AMP^R$ | This study |
| pAct2.2 Krr1 193-256 | *GAL4 AD-KRR1* 193-316 257stop *2µ LEU2* $AMP^R$ | This study |
| pAct2.2 Krr1 193-246 | *GAL4 AD-KRR1* 193-316 247stop *2µ LEU2* $AMP^R$ | This study |
| pAct2.2 Krr1 193-247 | *GAL4 AD-KRR1* 193-316 248stop *2µ LEU2* $AMP^R$ | This study |
| pAct2.2 Krr1 193-316 PFPP-GSGG | *GAL4 AD-KRR1* 193-316 P250G F251S P252G P253G *2µ LEU2* $AMP^R$ | This study |
| pAct2.2 Krr1 FL | *GAL4 AD-KRR1 2µ LEU2* $AMP^R$ | This study |
| pAct2.2 Krr1 1-316 PFPP-GSGG | *GAL4 AD-KRR1* P250G F251S P252G P253G *2µ LEU2* $AMP^R$ | This study |
| pAct2.2 RPS14A (uS11) | *GAL4 AD-uS11 2µ LEU2* $AMP^R$ | This study |
| pAct2.2 LargeT | *GAL4 AD-LARGET 2µ LEU2* $AMP^R$ | Dual Systems |
| pRS315 Krr1 WT | $P_{KRI1}$ *KRR1 CEN LEU2* $AMP^R$ | Twist Bioscience |
| pRS315 Krr1 1-247 | $P_{KRI1}$ *KRR1 1-247 CEN LEU2* $AMP^R$ | Twist Bioscience |
| pRS315 Krr1 1-266 | $P_{KRI1}$ *KRR1 1-266 CEN LEU2* $AMP^R$ | Twist Bioscience |
| pRS315 Krr1 PFPP-GSGG | $P_{KRI1}$ *KRR1 P250G F251S P252G P253G CEN LEU2* $AMP^R$ | Twist Bioscience |
| pRS315 Krr1 YTPFPP-SGGSGG | $P_{KRI1}$ *KRR1 Y248S T247G P250G F251S P252G P253G CEN LEU2* $AMP^R$ | Twist Bioscience |
| pRS315 Krr1 1-266 + YTPFPP-SGGSGG | $P_{KRI1}$ *KRR1 1-266 Y248S T247G P250G F251S P252G P253G CEN LEU2* $AMP^R$ | Twist Bioscience |
| pRS313 Krr1-TAP | $P_{KRI1}$*KRR1-3xFLAG-TEV-ProtA CEN HIS3* $AMP^R$ | Twist Bioscience |
| pRS315 Fap7 WT | $P_{FAP7}$ *FAP7 CEN LEU2* $AMP^R$ | This study |
| pNopGFP1L-Fap7 | *GFP-FAP7 CEN LEU2* $AMP^R$ | This study |
| pRS315 *Si*Fap7 | $P_{FAP7}$ *SiFAP7 CEN LEU2* $AMP^R$ | Twist Bioscience |
| pRS315 *Si*Fap7-GFP | $P_{FAP7}$ *SiFAP7-GFP CEN LEU2* $AMP^R$ | Twist Bioscience |
| pRS315 CtFap7 | $P_{FAP7}$ *CtFAP7 CEN LEU2* $AMP^R$ | Twist Bioscience |
| pRS315 TbFap7 | $P_{FAP7}$ *TbFAP7 CEN LEU2* $AMP^R$ | Twist Bioscience |
| pRS425 TbFap7 | $P_{FAP7}$ *TbFAP7 2µ LEU2* $AMP^R$ | Twist Bioscience |
| Yep315 $P_{GAL1}$ Fap7 WT | $P_{GAL1}$ *FAP7 2µ* | This study |

| | | |
|---|---|---|
| Yep315 $P_{GAL1}$ SiFap7 WT | *$P_{GAL1}$ SiFAP7 2µ* | This study |
| pGEX-6P-1 | GST *$AMP^R$* | Demmel et al. (2008)[4] |
| pGEX-6P-1 Fap7 | GST-Fap7 *$AMP^R$* | Peña et al. (2016)[1] |
| pGEX-6P-1 Tsr4 | GST-Tsr4 *$AMP^R$* | Oborská-Oplová et al. (2025)[5] |
| pGEX-6P-1 *Ta*Fap7 | GST-TaFap7 *$AMP^R$* | GenScript |
| pET47b $His_6$-Fap7 | $His_6$-Fap7 *$KAN^R$* | Peña et al. (2016)[1] |
| pET47b $His_6$-Krr1 | $His_6$-3C-Krr1 *$KAN^R$* | This study |
| pET47b $His_6$-Krr1 CTT | $His_6$-Krr1 193-316 *$KAN^R$* | This study |
| pGEX-6P-1 Fap7:uS11 | GST-Fap7 uS11 *$AMP^R$* | Peña et al. (2016)[1] |
| pETCola-1 *Si*Fap7 uS11 | $His_6$-SiFap7 uS11 *$KAN^R$* | Twist Bioscience |
| pEt29b GST-Kri1 PEP | GST-Kri1 51-96 209-267 *$KAN^R$* | Twist Bioscience |
| pCOLADuet-1 fap7-2:uS11 | $His_6$-Fap7 D82A H84A:uS11 *$KAN^R$* | Peña et al. (2016)[1] |
| pEM1 Krr1 | $His_6$-GB1-TEV-Krr1 *$AMP^R$* | This study |

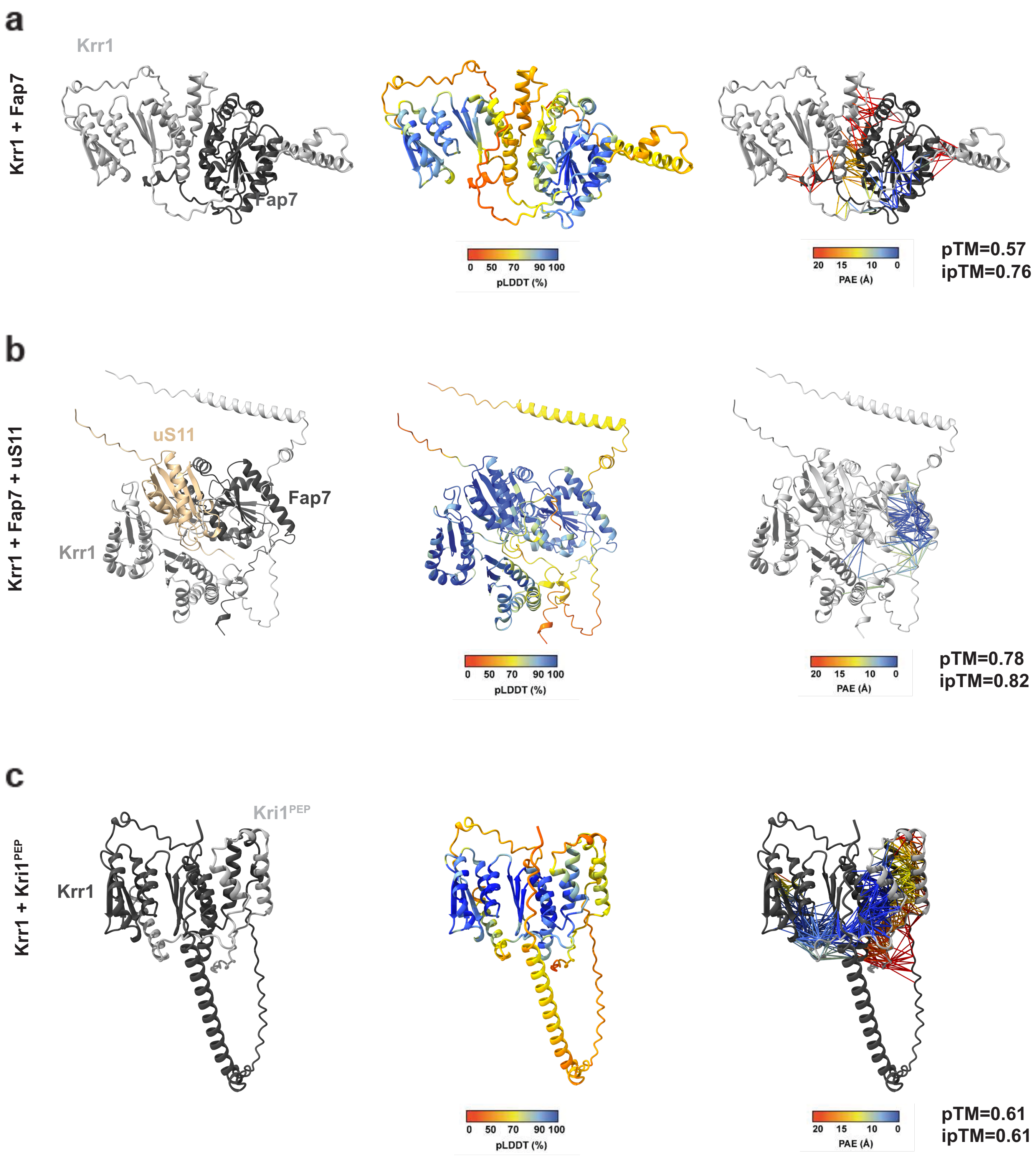


**Supplementary Figure 1. Structural AlphaFold models of Krr1-containing complexes. a** AlphaFold2-derived models of Krr1 (light grey) binding to Fap7 (dark grey). **b** AlphaFold2-derived models of Krr1 (light grey) in complex with Fap7 (dark grey) and uS11 (beige). **c** AlphaFold2-derived models of Krr1 (dark grey) binding to Kri1 peptide (light grey). **(a-c)** Predicted local distance difference test (pLDDT) and predicted aligned error (PAE) scores for the indicated AlphaFold2-predicted complexes. The pseudobonds indicating the PAE between pairs of residues were set for 8 Å. Structures were analyzed using ChimeraX (version 1.7.1, UCSF Chimera, National Institutes of Health, CA, USA)[6].

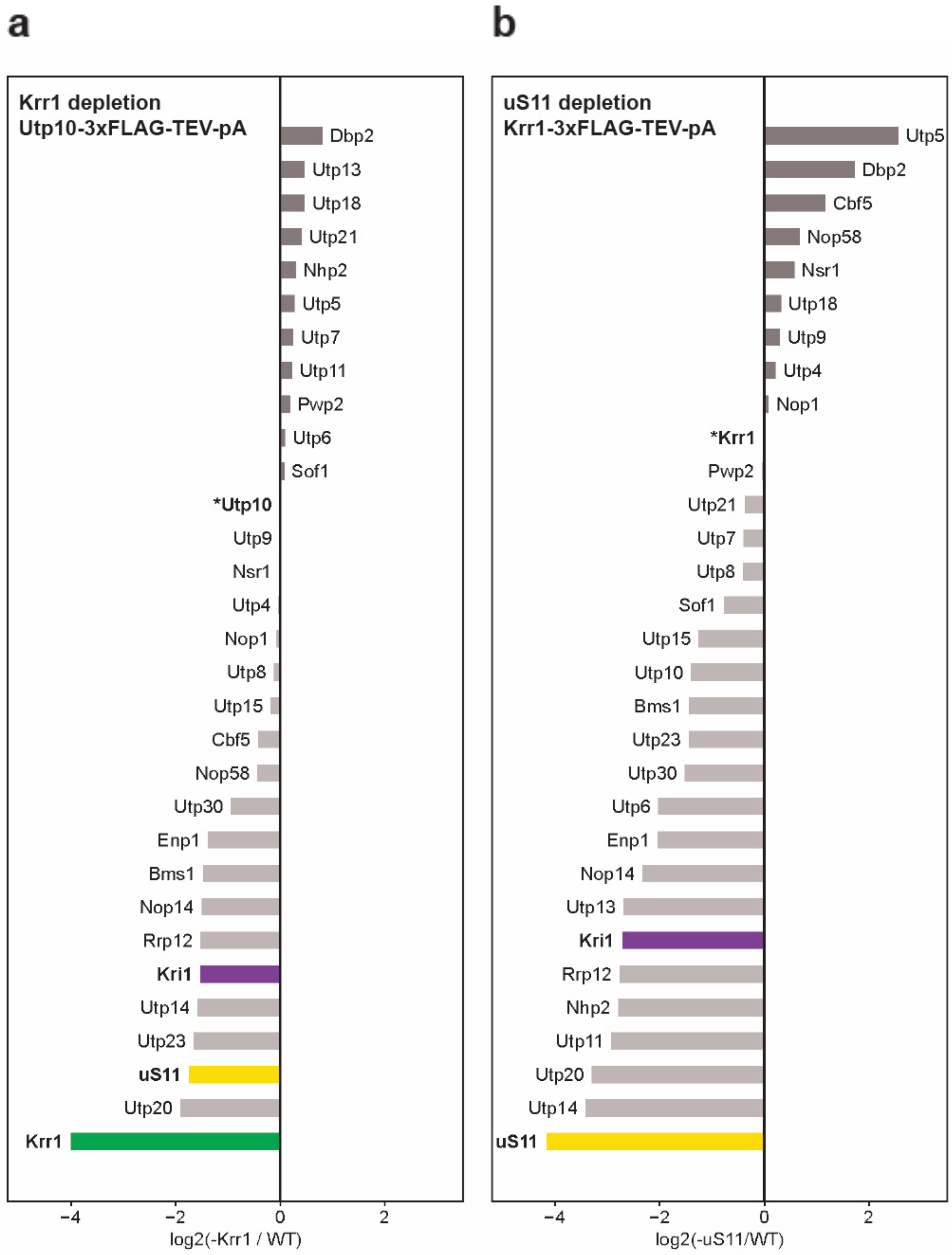


**Supplementary Figure 2. Semiquantitative mass spectrometry analysis of Krr1-TAP and Utp10-TAP purifications upon depletion of 90S biogenesis factors. a** Utp10-TAP was affinity-purified from cells grown under wild-type conditions or upon Krr1 depletion. Co-enriching assembly factors were quantified by mass spectrometry, and a subset of the factors are shown as log2 fold enrichment in -Krr1 relative to WT. Values were normalized to the Utp10 bait protein signal[5]. **b** Krr1-TAP was affinity-purified from cells grown under wild-type conditions or upon uS11 depletion. Co-enriching assembly factors were quantified by mass spectrometry, and a subset of the factors are shown as log2 fold enrichment in -uS11 relative to WT. Values were normalized to the Krr1 bait protein signal. In both panels, bait proteins are indicated by an asterisk. Krr1, uS11, and Kri1 are highlighted in green, yellow, and purple, respectively[7].

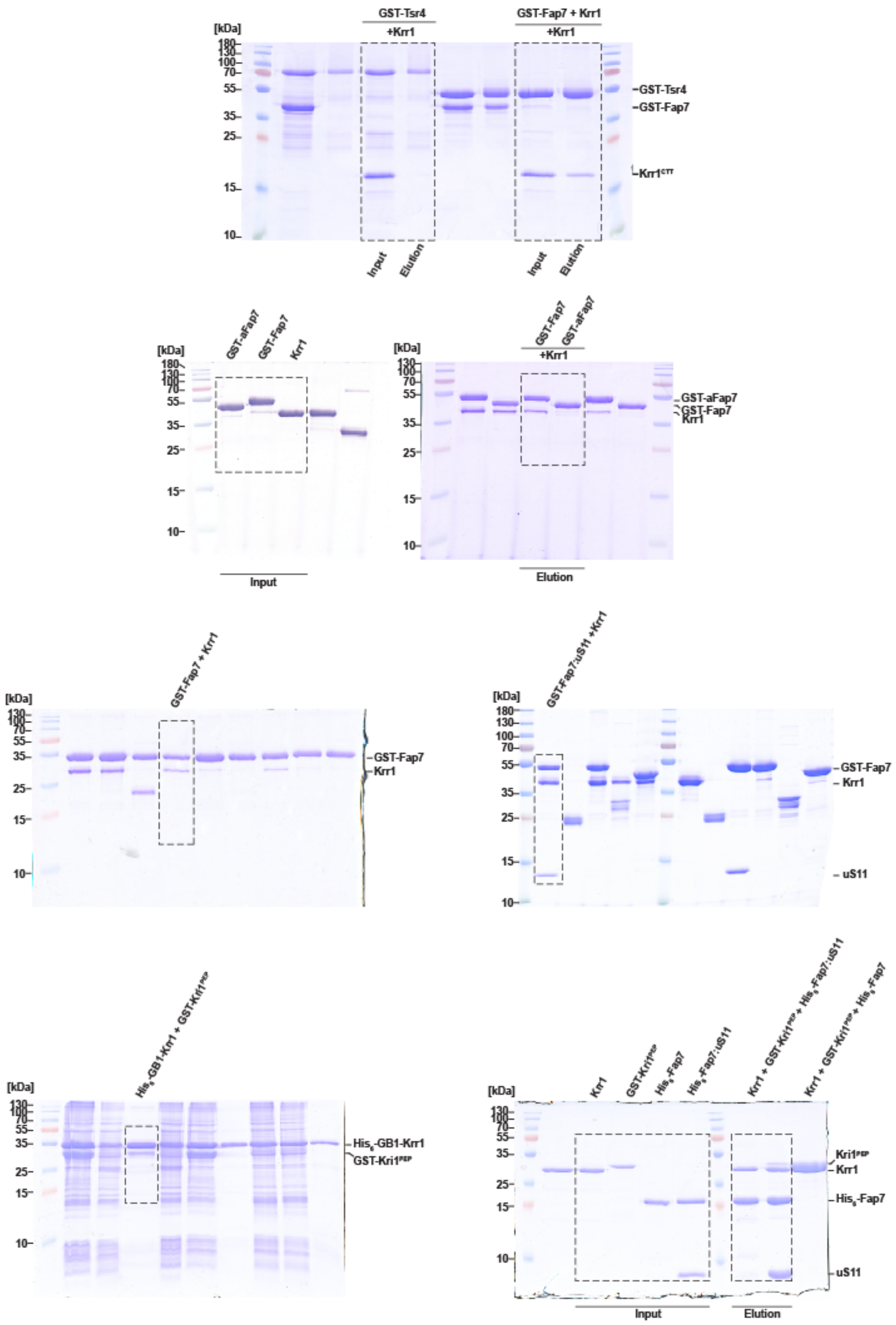


**Uncropped SDS-PAGE Gels for Fig. 3b; Fig. 4d; Figs. 6a, left panels; Fig. 6d.**
